# Discovery of a Novel 3site State as the Multi-Substrate Bound State of P450cam

**DOI:** 10.1101/2023.03.15.532864

**Authors:** Mohammad Sahil, Tejender Singh, Soumya Ghosh, Jagannath Mondal

**Affiliations:** Tata Institute of Fundamental Research, Hyderabad 500046, India

## Abstract

Archetypal metalloenzyme Cytochrome P450cam (CYP101A1) catalyzes regioselective hydroxylation of its native substrate camphor in heme active site. However, the proposal of potential existence of additional substrate binding modes distal from the active site in P450cam and their concomitant roles in regulating recognition at active site have remained a matter of recurring discourse. Herein we report the discovery of a novel *3site* state in P450cam, where three substrate molecules were observed to simultaneously bind to P450cam at three distinct sites including the heme active site. These three binding modes, hereby referred as *catalytic*, *waiting* and *allosteric* binding modes in *3site* state, are allosterically inter-linked and function in mutually synergistic fashion. The *3site* state possesses regio-selective conformations of substrate essential for catalysis and establishes substrate-ingress and product exit process to and from the active site via two distinct channels. The ensemble of three-state binding modes are found to be self-consistent with NMR pseudo-contact shift data obtained from TROSY-HSQC measurements and DEER based predictions. Binding of redox partner Putidaredoxin with *3site* model retains closed conformation of 3site state, siding with NMR based hypothesis that the catalysis would take place in closed insulation of P450cam even in presence of its redox partner.

**Statement of Significance:** Ubiquitous superfamily of mono-oxygenases cytochrome P450s are involved in broad range of metabolic process in all domains of life and are also important drug targets. Apart from the well known and established binding mode in heme active site, the substrate bindings at additional distal sites have been postulated in multitude of P450s. Using the archetypal bacterial cytochrome P450 i.e., P450cam, a novel *3site* state of cytochrome P450 is elucidated in this work. The novel 3site state has two additional binding modes namely *waiting* and *allosteric* (also postulated previously), apart from known binding mode *catalytic* in the active site. The known functions of P450cam are found to be most optimally explained by this 3site state, instead of single substrate bound catalytic state. This state can be of critical importance for CYP superfamily at large and potentially be useful in understanding the non-Michaelis behaviour, observed in many P450s.

## Introduction

Cytochrome P450s (CYPs), one of the largest enzyme families, are b-type heme containing mono-oxygenases that are involved in broad range of metabolic processes such as drug detoxification, steroid biosynthesis, fatty acid metabolism etc.^1^ The ubiquitous CYP super-family catalyses the stereospecific and regiospecific hydroxylation of aliphatic and aromatic substrates using molecular oxygen in all three domains of life.^2^ The overall P450 fold and heme-chemistry is conserved from bacterial P450s to higher mammals such as humans.^1^ In humans, 57 P450s have been identified till date and has been characterized as important pharmaceutical targets which are responsible for metabolising around half of currently used drugs.^3^ The archetypal P450 CYP101A1 (popularly known as P450cam) was the first cytochrome P450 to be sequenced, characterized and one of the most thoroughly investigated.^4,5^ It has served as the model P450 by contributing significantly to what is known of CYP superfamily functioning till date. ^6^

P450cam is constituted of heme active site i.e., the reaction center and an opening named channel-1 that link the completely buried active site with exterior of the enzyme ^7,8^ (Figure 1A). The channel-1 serves as the binding route through which substrate ingress to heme active site and gets hydroxylated.^9^ Channel-1 samples substrate dependent open and closed conformational extremes such that it explores open conformation prior to substrate binding and transitions to closed conformation upon substrate binding. ^10^ Through channel-1, P450cam recognises D-Camphor (referred in literatures as ‘cam’) as substrate among others and perform its hydroxylation to 5-exo-hydroxy-camphor (Figure 1B). In majority of the investigations since its first characterisation, only one substrate binding mode was known for P450cam i.e., in the heme active site, to which the substrate dependent closing of channel-1 and other substrate dependent conformational changes have been attributed. Albeit, over decades, there have been multiple affirmations that, apart from the well-established binding mode at the heme active site, there might exist additional substrate-binding modes in P450cam or CYP superfamily at large.

**Figure 1:**
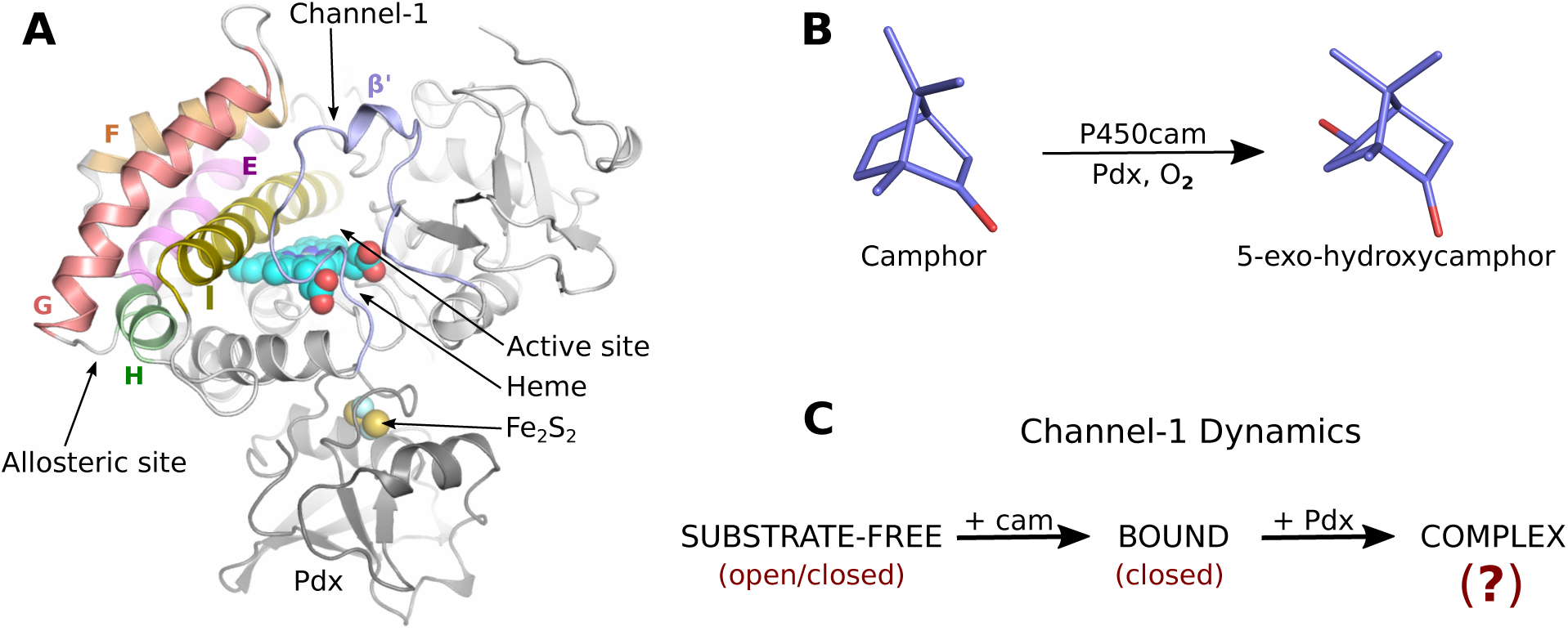
The cytochrome P450cam: (A) Crystal structure of cytochrome P450cam-Pdx complex i.e., CYP101A1 from *Pseudomonas putida* (PDB id - 4JX1). Important secondary structures, active and putative allosteric site and channel-1 are highlighted. (B) Camphor hydroxylation to 5-exo-hydroxycamphor by P450cam in presence of Pdx and *O*_2_. (C) Channel-1 dynamics in P450cam in presence/absence of substrate camphor (cam) and Pdx. State of channel-1 in P450cam-Pdx (complex) is controversial at present.

As the first instance, based on NMR paramagnetic *T*_1_ relaxation, camphor was proposed to bind to an additional site which was estimated to be 15-16 °*A* away from active site. ^11^ It had been also proposed for P450cam and other CYPs that substrate might undertake multi-step binding process such that it might bind an additional site before actually transitioning to active site.^12^ Hence, the NMR posited second weak binding site was postulated to be the first step of either multi-step or induced fit binding process.^13^ Recently, an additional binding mode near the junction of E, F, G and H helices (allosteric site, Figure 1A) was posited to be the second binding mode of P450cam that had remained hidden until recently.^14^ The allosteric site was proposed to be involved in substrate dependent allostery such that it facilitates active site binding by opening the channel-1. The newly posited allosteric site was found to be conserved across several members of CYP superfamily. ^15^ However, substrate binding at allosteric site further complicates the channel-1 dynamics. Current standing on this field concludes that two camphor bind to P450cam in the active and allosteric sites. ^16^ However binding at the former (active site) closes the channel-1, while binding at the latter (allosteric site) is proposed to open the channel-1.^16^ This proposal is counterintuitive as the opposing effect of two camphor molecules on channel-1 is in direct disagreement with known substrate dependent closing of channel-1.^10^ Additionally, in several other important CYPs, multiple binding modes both within and outside the active site have often been proposed through diverse methodologies.^17–22^ Despite multiple direct/indirect evidences, any such state of cytochrome P450 and its functioning have not yet been established.

The controversial effector role of redox partner putidaredoxin (Pdx) further intermingled with presence of multiple binding modes in P450cam. Pdx binds P450cam on distal side (opposite to channel-1, Figure-1A) and contributes two electrons to heme active site, necessary for hydroxylation reaction. It was believed that the hydroxylation reaction occurs in closed conformation of channel-1 (substrate induced), until redox partner Pdx was postulated crystallographically to open back the channel-1.^23^ The Pdx induced opening of P450cam was challenged shortly after its publication and crystal and NMR structure of P450cam-Pdx complex were determined in intermediate and closed state respectively. ^24^ At present, the channel-1 conformation in presence of Pdx stands controversial^25^ with Pdx induced opening supported by DEER^26^ and MD^27,28^ observations in contrast to NMR Pseudo contact shift (PCS) data^29^ and other biochemical analysis^30^ (Figure 1C). It has been postulated indirectly that additional binding mode in P450cam (allosteric site) can function in synergy with Pdx in facilitating product egress from the heme active site. ^14^ Recently, while the effect of multiple substrate and Pdx binding in P450cam’s product egress has been claimed to observed crystallographically, the conspicuous absence of the additional binding modes was very apparent.^31^ Overall, the postulated state of P450cam in presence of multiple substrates and redox partner Pdx has not been characterised yet.

The present work is an effort to resolve the fuzzy picture around highly debated existence of additional binding modes of P450cam and conformation of channel-1 in absence and presence of redox partner Pdx. In particular, pertinent to this topic, here following questions were probed:

- What are the possible distinct substrate-binding sites around P450cam?
- How do these binding modes mutually communicate with each other in P450cam?
- How does presence of additional binding modes control dynamics of entry channels in absence or presence of redox partner pdx ?

Towards this end, we have undertaken an initiative of extensive computer simulation to exhaustively explore the potential substrate-binding modes in P450cam. As would be revealed in the following section, as a novel observation, three distinct substrate binding modes in P450cam were reproducibly observed. The novel three substrate bound conformation (referred here as ‘*3site state*’) is hereby proposed to be the eventual bound state of P450cam. For validation, the hereby claimed state was shown to reconcile precedent NMR, crystallographic and other measurements of P450cam. Interestingly, it was found that known multifaceted aspects of P450cam including the controversial role of Pdx can only be explained in light of hereby proposed state and not by traditionally known substrate-bound state in active site. This work provides comprehensive and detailed structural view of both the aspects of P450cam i.e., unknown binding modes and role of pdx in substrate recognition.

## Results

### 1: A *3site state* represents the final bound state in P450cam

We started our investigation by simulating the substrate binding modes in P450cam using unbiased binding simulations,^32–34^ where ligand molecules unbiasedly search for its binding mode(s). In this regard, multiple long unbiased binding simulations were initiated by randomly placing four copies of camphor (substrate in this work) molecules around substrate-free P450cam (open conformation) in the solvent. Contrary to precedent investigations, ^14^ the substrate in our initial configuration had no physical contact with the receptor, hence the substrate had no a-priori knowledge of any binding site. During the course of the simulations the substrate explored diverse locations around the protein in an unbiased manner, searching for binding modes at various location. As a novel observation, three camphor molecules eventually settled at three distinct binding sites of P450cam (Figure 2A,B). The spatial density map of the camphor showed three large distinct densities corresponding to three binding modes representing the most explored regions of P450cam (Figure 2B). The three binding modes are hereafter named as (Figure 2A,B, S1A-C):

1. ***catalytic*** - the traditional and previously known binding mode present at the catalytic center: in this mode camphor binds just above the heme moiety and hydrogen bonded to Y96 with a *k_on_* of 5.5 × 10^7^ *M^−^*^1^*s^−^*^1^. This is the site of reaction where catalytic substrate undergo hydroxylation;
2. ***waiting*** - additional camphor molecule binds along channel-1 in the active site but significantly above catalytic binding mode with a *k_on_* of 9.9 × 10^7^ *M^−^*^1^*s^−^*^1^, in this binding mode camphor waits for its turn to be catalyzed;
3. ***allosteric*** - one camphor molecule binds at the previously proposed allosteric site^14^ ( at the junction of E, F, G and H helices and constituted by V123, L127, L166, T217 etc) with *k_on_* of 8.1 × 10^8^ *M^−^*^1^*s^−^*^1^ and hydrogen bonded to T217.

**Figure 2:**
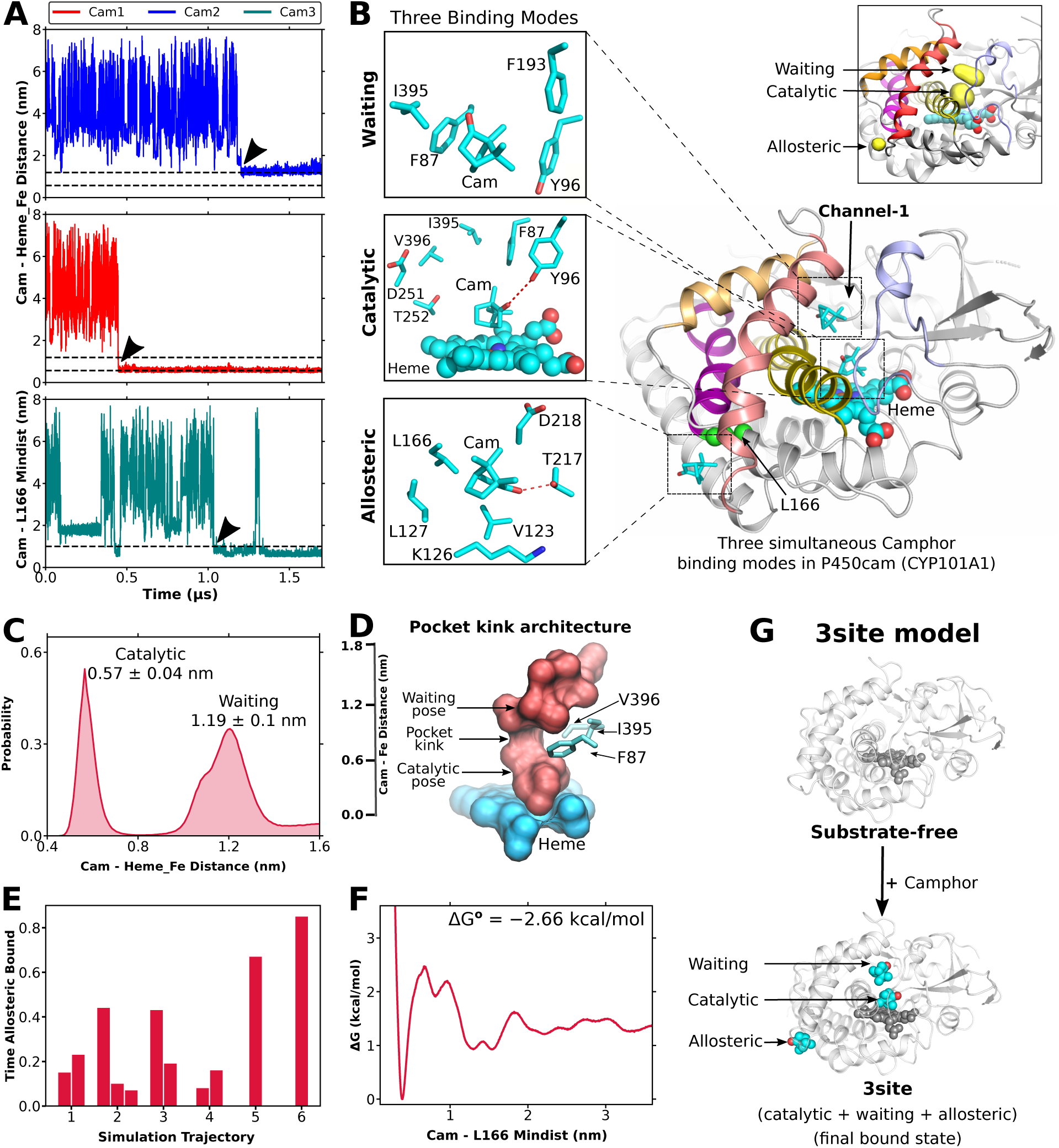
Novel 3site state in P450cam. (A) Simultaneous binding of three substrate molecules to P450cam at *catalytic* (cam1), *waiting* (cam2) and *allosteric* (cam3) binding modes for one of the binding simulations. (B) Three camphor binding modes in P450cam. Inset: The spatial density map of substrate in binding simulation showing three densities corresponding to three binding modes. (C) Probability distribution of camphor localization with respect to Heme-Fe showing discrete catalytic and waiting modes within the channel-1. (D) Pocket kink within the channel-1 of P450cam, dissecting the active site for distinct catalytic and waiting modes. Crimson surface represents the shape of the binding pocket and scale bar in accordance with subfigure-C. (E) Fraction of total time a camphor remain bound to allosteric site. Each bar represents a camphor molecule. (F) Binding energy of allosteric mode measured by camphor probability distribution in binding simulations. (G) Proposed *3site* model of substrate recognition in P450cam.

The binding poses in the metal-active-site i.e., catalytic and waiting modes transpired through previously posited^9^ channel-1 (Figure 1A, S1D), while allosteric site represents a surface exposed cavity and direct binding of the substrate from the solvent to this site was observed.

While the catalytic binding mode represents the previously known and well established binding mode (Figure S1E), we needed to assess the stability and reproducibility of waiting and allosteric binding modes. The P450cam active site in open conformation is large enough with a volume of 1136.9 *A*^3^ and can accommodate two camphor molecules (space-filling volume 160 *A*^3^). Within the active site, the catalytic and waiting modes represent discrete and mutually separated binding modes such that the substrate in ‘catalytic mode’ binds within a distance of 0.57 ± 0.04 nm from Heme-Fe while the substrate in ‘waiting mode’ binds at 1.19 ± 0.1 nm away from Heme-Fe. (Figure 2C). The pocket architecture indicates the presence of a bottleneck (hereafter named as ‘pocket-kink’) constituted by F87, I395 and V396 which keep the catalytic and waiting modes distinctly coexisting inside the channel-1 leading to active site (Figure 2D). On the other hand, the allosteric site represents the high fidelity cryptic pocket and had also been postulated previously.^14^ Statistically one or more camphor binding to the allosteric site was observed in all binding simulations (Figure 2E, S1F-H). Revisiting our previous binding simulations^9^ also revealed allosteric site bindings in every simulations (Figure S1G). Overall, the allosteric mode represents a stable binding mode with Δ*G°* = -2.66 kcal/mol (Figure 2F). Above all, the simultaneous binding of three substrate molecules to P450cam was reproducible across multiple independent replicates within few microseconds of real time (Figure S2). On the basis of above findings, we propose a new model of substrate recognition in P450cam, hereafter named as *3site model* (Figure 2G). The 3site model proposes that instead of traditionally known bound state (only catalytic binding mode), the three substrate molecules simultaneously bind to P450cam and represent the thermodynamically most stable bound state (hereafter named as 3site state). The novel 3site state is distinct from previously postulated multi-substrate bound states^14^ where substrates bound at catalytic and allosteric sites. Additionally as explained in incoming sections, simultaneous bindings at all three binding modes is crucial for P450cam to be functional, dictating the uniqueness of 3site state.

To understand the distinct roles played by three binding modes with better statistics, individual structural ensembles of each possible combinations of single and doubly bound states leading to the proposed 3site model were generated. All potentially possible states in 3site model and nomenclature used are represented in Figure 2A-F. For example, the ‘catalytic state’ represents P450cam with substrate camphor in catalytic binding mode. Hereafter catalytic mode refers to camphor bound to catalytic binding mode while catalytic state refers to simulation ensemble of P450cam with camphor bound to catalytic mode; and similarly for other states. For each state, P450cam in open conformation with substrate molecule(s) in desired binding modes was taken as the starting structure. The starting structures were generated by placing the camphor molecules in the desired binding modes based on the knowledge from aforementioned unbiased binding simulations. To generate the ensemble, at least three independent simulations were performed to a maximum of one microsecond. The system was allowed to take its favourable conformation(s) for a particular amount of time in unbiased simulation. An analysis based on the RMSD of protein indicates that it took around 350 nanosecond (ns) for any of these states to acquire the state specific conformational changes (Figure S3), hence simulation trajectories after 350 ns were considered for analysis. In this way, all the states were simulated from same ‘open conformation’ but generated different ensembles where each state’s ensemble possessed the conformations specific to that particular state. Ultimately, each state represents a large ensemble of approximately 1.95×10^5^ conformations. Note worthy, as explained in the upcoming paragraph, the waiting state is not stable by itself and only stable in 3site state, hence its solo ensemble (i.e. in absence of the substrates in other two binding modes) could not be generated.

### 2.1: Binding modes in 3site state are allosterically linked

To be functional, the three binding modes of the 3site state should be allosterically linked with each other or at the minimum, the waiting and allosteric modes should be able to allosterically impact the catalytic mode (the substrate that will undergo hydroxylation). To confirm the allosteric linkage between different binding modes, the ‘principle of allosteric reciprocity’ was investigated. ^35,36^ Hence, we assessed the stability of the substrate in a particular binding mode by comparing their residence times or free energetics with/without the presence of substrate(s) in distal binding mode(s). For allosteric mode, the residence time^37^ of substrate camphor in allosteric pocket was calculated as a measure of its stability, using infrequent metadynamics simulation approach.^38^ Unbinding times from 50 independent metadynamics replicas were calculated and residence time was estimated by fitting the unbinding times with poisson cumulative probability distribution (methods). With active site devoid of its binding modes i.e., unbound, the substrate in the allosteric mode resides for 4.1 ± 0.2 *μ*s of residence time (Figure 3G, Table-1). With the substrates bound to catalytic and/or waiting binding modes, the residence time of camphor in allosteric site significantly increased to 14.1 ± 0.7 *μ*s (Figure 3G, S4A,B, Table-1). For catalytic binding mode, the standard free energy of camphor binding to active site (at catalytic mode) was calculated as measure of its stability using umbrella sampling approach (methods). With allosteric site unbound, the catalytic mode recognise the substrate with a stability of Δ*G°* -5.4 ± 0.3 kcal/mol. However, the stability of the substrate bound to catalytic mode significantly increases with Δ*G°* of -7.6 ± 0.6 kcal/mol when the allosteric site was also occupied by substrate (Figure 3H, Table-1). Catalytic mode being the previously reported binding mode, its binding free energy, as derived from *k_D_*, had been previously estimated^30^ with Δ*G°* of -8.1 kcal/mol, which can only be explained in presence of substrate at the allosteric mode. This clearly indicates that the camphor binds to both active and allosteric sites. On the other hand, the ‘waiting binding mode’, as mentioned in the aforementioned paragraph, was found to be unstable by itself i.e., in waiting state it either transitions to catalytic mode or unbinds and moves back to solvent (Figure S4C). The waiting mode was found to be completely stable only in 3site state (Figure S4C). Waiting mode was estimated to have a residence time of 0.6 ± 0.08 ms without allosteric mode, but a very stable residence time of 644.9 ± 155.8 ms in 3site state (Figure 3I, S4D, Table-1). The overall trend of stability suggests that all three binding modes in 3site state allosterically reinforces each other’s stability. The overall allosteric stabilization of three binding modes occurred through G248 and G249 induced kink in I helix as explained in Supplementary results (SR-1).

**Table 1:** Binding free energies and residence times of three binding modes in different states of P450cam

|  |  |  |
| --- | --- | --- |
| **Catalytic**<br>Binding energy (kcal/mol) | -Allosteric<br>+Allosteric | $-5.4 \pm 0.3$<br>$-7.9 \pm 0.6$ |
| **Catalytic**<br>Residence time (ms) | Channel-1 (+Allosteric)<br>Channel-1 (-Allosteric)<br>Channel-2 (3site) | $1.4 \pm 0.2 \times 10^7$<br>$4.6 \pm 1.0 \times 10^3$<br>$9.8 \pm 0.7 \times 10^1$ |
| **Waiting**<br>Residence time (ms) | -Allosteric<br>+Allosteric | $0.6 \pm 0.08$<br>$644.9 \pm 155.8$ |
| **Allosteric**<br>Residence time (μs) | -Active<br>+Active (catalytic)<br>+Active (catalytic+Waiting)<br>-Active (L166A)<br>-Active (T217V) | $4.1 \pm 0.2$<br>$11.4 \pm 1.0$<br>$14.1 \pm 0.7$<br>$0.166 \pm 0.003$<br>$16.1 \pm 0.7$ |

**Figure 3:**
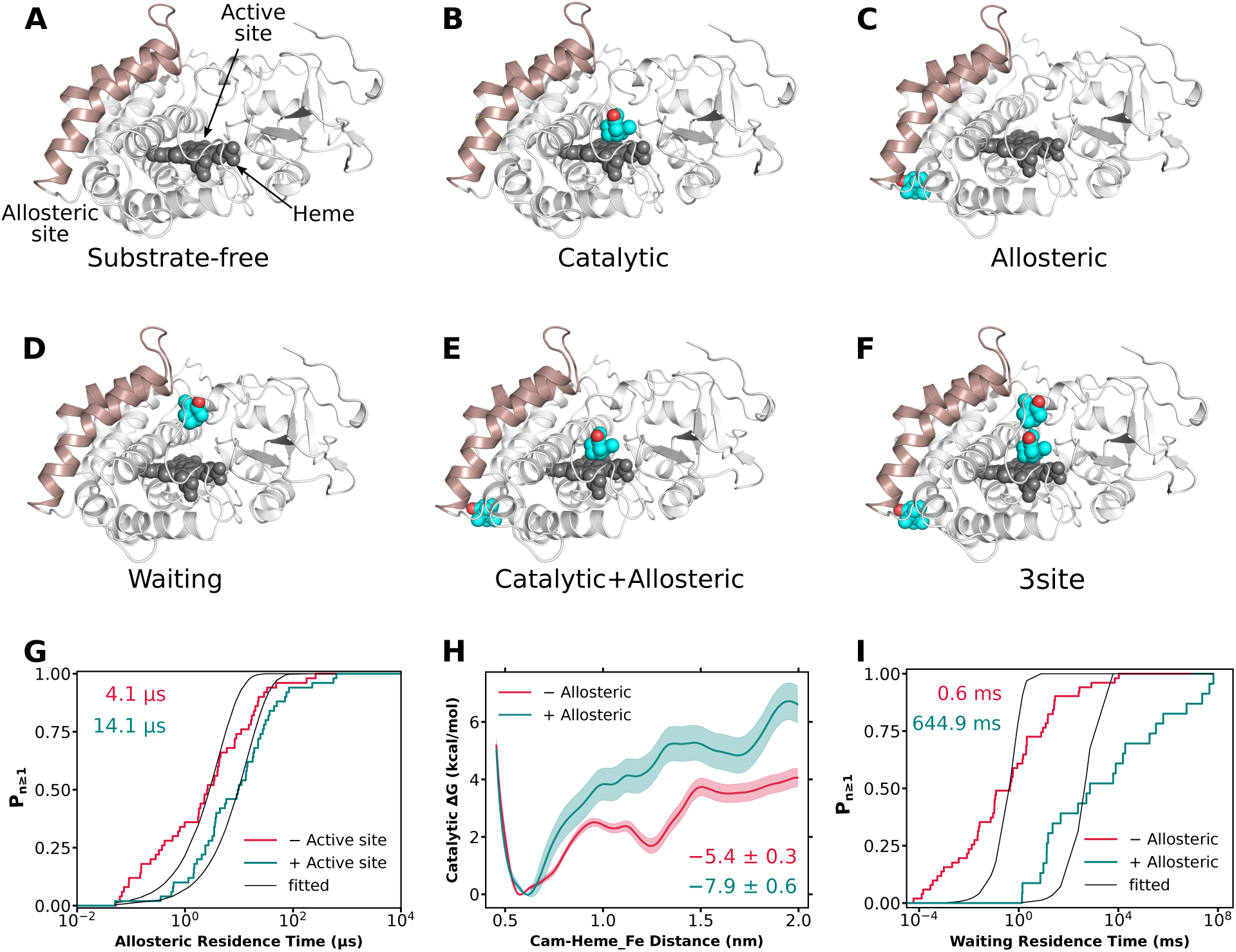
Allosteric stabilization of three binding modes: (A-F) Starting configurations used to generate the simulation ensembles of possible states of 3site model, labelled with nomenclature used in this work. Also see Figure S13. (G) Probability of allosteric mode unbinding with and without active site binding modes i.e., both catalytic and waiting modes. Values on top represent the residence times of allosteric mode. Goodness of fit is 0.27 (−Active site) and 0.4 (+Active site). (G) Binding free energy of catalytic mode with and without the allosteric mode. Width of curve represents the error. Values represents the standard binding free energy. (H) Probability of Waiting mode unbinding with and without the allosteric mode. Values represents the residence times of waiting mode. Goodness of fit is 0.013 (−Allosteric) and 0.025 (+Allosteric).

### 2.2: Binding modes in 3site state function synergistically

The observations that three binding modes of 3site state represent the final bound state of P450cam and stabilise each other lead to an interesting question: whether 3site binding modes function together i.e., whether the binding associated conformational changes in P450cam are induced by only one of the binding modes or by all three binding modes together. In this regard, we investigated three particular conformational changes in the active site that are known to be crucial for function i.e., a) Y96 conformation, b) its hydrogen bond with catalytic mode, and c) pocket kink. As a key hallmark of substrate recognition in P450cam, the substrate camphor forms hydrogen bond with Y96 side-chain in active site. ^14,39^ Y96 was found to exist in two conformations as measured by its *β* dihedral N-C*α*-C*β*-C*γ*, *conf-a* (−70*°*) and *conf-b* (−170*°*) (Figure 4A). In substrate-free state, Y96 predominantly exists in *conf-a*, pointing away from catalytic center and hence not in a suitable conformation to form hydrogen bond with catalytic mode. Ideally, camphor binding should induce *conf-a* to *conf-b* transition of Y96, leading to the formation of Y96-camphor hydrogen bond. Interestingly, substrate-binding in the catalytic mode shifted the Y96 conformation from *conf-a* to *conf-b* but partially such that both conformations are equally likely in catalytic state. Additional binding of substrate in allosteric mode further shifts Y96 in favour of *conf-b*. Finally Y96 adopted *conf-b* predominantly in 3site state when all three binding modes are occupied by the substrate (Figure 4B). The results indicate that the transition of Y96 from *conf-a* to *conf-b* follows a ‘conformational selection’ ^40^ model of allostery, but interestingly *conf-b* was found to be synergistically stabilized together by all three binding modes. The combined effect of three binding modes synergistically produced conformational change of Y96 conformation in active site. Identical to Y96 conformation, Y96-camphor hydrogen bond formed stably only in 3site state by a synergistic effect of three binding modes (Figure S5).

**Figure 4:**
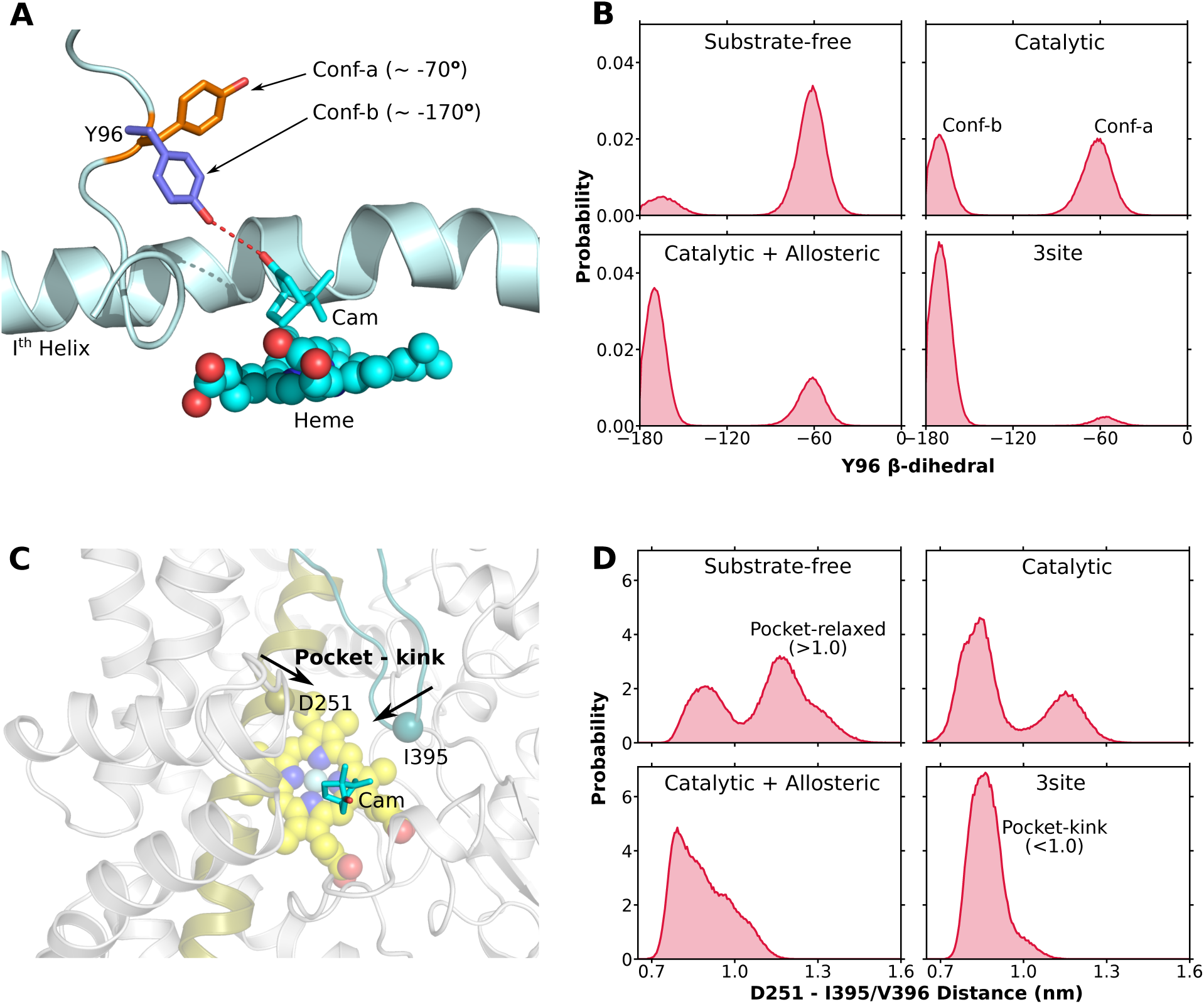
Synergistic functioning of three binding modes: (A) Two conformations of Y96 sidechain. (B) Y96 conformation in different states of P450cam. (C) Pocket architecture around catalytic mode. Arrows indicate motions contributing to change in pocket architecture in active site, i.e., pocket-kink. (D) Pocket-kink in different states of P450cam as measure by distance between Ca atoms of D251 and I395/V396.

Noteworthy, Y96 though important, is not the sole parameter which holds on the active site binding mode, as Y96F mutant does not change the regio-selectivity leading to the idea that pocket architecture also holds on to the catalytic camphor and necessary for function.^39^ As discussed in previous section, the active site was found to posses a ‘pocket kink architecture’ which explained the existence of discrete waiting and catalytic modes within the active site (Figure 2D). The position of C-terminal region (residues 390-400) of P450cam with respect to I helix generates the pocket kink within the active site (Figure 4C). In substrate-free state, active site mainly exists in the relaxed state with D251-I395/V396 distance more than 1 nm (Figure 4D). Upon binding in catalytic mode, the formation of pocket kink started but not completely. Monotonically and synergistically a predominant pocket kink was generated only in 3site state (Figure 4D). Together, these reconstruction-based analysis indicates that conformational changes in P450cam were acquired gradually from substrate-free to 3site state and was mediated by consecutive binding of substrates in three binding modes. In other words, three binding modes function synergistically and together induce conformational changes in P450cam.

### 3.1 : 3site state reconciles with NMR and DEER measurements

The finding of a novel 3site state necessitates its validation/agreement with previously reported observations. Here reported 3site model concludes that three substrate molecules shall simultaneously bind to P450cam and all three binding modes are necessary for function as these are allosterically linked and function synergistically (Figure 2-4). In such a scenario, the 3site state, instead of catalytic state, should predominantly exist in solution in presence of substrate camphor. Towards this end, we checked for agreement between the simulation ensembles of 3site model and previously reported measurements.

Firstly, the pseudo-contact shift (PCS) data, generated via NMR measurements for various states of P450cam, were compared with the PCS calculated from simulation ensembles. PCS is the difference between chemical shifts in paramagnetically and diamagnetically tagged protein and represents a measure of state of protein in solution that can be fitted with structures at hand.^41^ Hiruma et. al and Skinner et. al have measured the ^15^*N* −^1^ *H* PCS of P450cam in solution both with and without substrate camphor. ^24,29^ The distance between the paramagnetic tag (CLaNP-7-Ln) and C*α* atoms of its attachment sites (E195, A199) on protein was calculated as part of fitting process. This C*α*-Ln distance was expected to fall within 7-9 °*A*^29,42^ and is used as an indicative of goodness of fit (methods). For instance, the substrate-free simulation ensemble fitted reasonably with ^1^*H^N^* Leu PCS of substrate-free P450cam with average C*α*-Ln distance of 9.5 °*A* (E195) and 7.4 (A199) °*A* and PCS from simulation and NMR are in good agreement (Figure S6A,B). In contrast, the substrate-free simulation ensemble does not fit with any PCS data observed in presence of camphor (Figure S6C).

These validations indicate that PCS data from solution NMR can be used to distinctly correlate the present computer simulated samples of P450cam. Therefore, two TROSY-HSQC derived PCS datasets measured in presence of substrate camphor were taken, namely (a) ^15^*N* −^1^ *H* PCS of P450cam residues^24^ (^1^*H^N^*), and (b) ^15^*N* −^1^ *H* PCS of leucine residues of P450cam^29^ (^1^*H^N^* Leu). The solution NMR observed PCS datasets were fitted with catalytic and 3site simulation ensembles, to understand which state predominantly exist in solution. Interestingly, the catalytic state did not fit well with ^1^*H^N^* PCS data with average C*α*-Ln distances of 6.78 and 11.1 °*A* for E195 and A199 respectively (Figure 5A). However on the other hand, the 3site simulation ensembles perfectly fitted with ^1^*H^N^* PCS data with average C*α*-Ln distances of 7.68 and 7.64 °*A* for E195 and A199 respectively (Figure 5B). The calculated PCS from 3site simulation ensemble also perfectly matched with NMR observed ^1^*H^N^* PCS with a very low Q-score^43^ of 0.045 (Figure 5C). Exactly same conclusions were also concluded for Leu ^1^*H^N^* PCS data which perfectly fitted with simulation ensemble of 3site state while not with the catalytic state (Figure S6D-F). Overall, these results indicated that instead of traditionally known catalytic state, 3site state predominantly exist as a final bound state of P450cam in solution.

**Figure 5:**
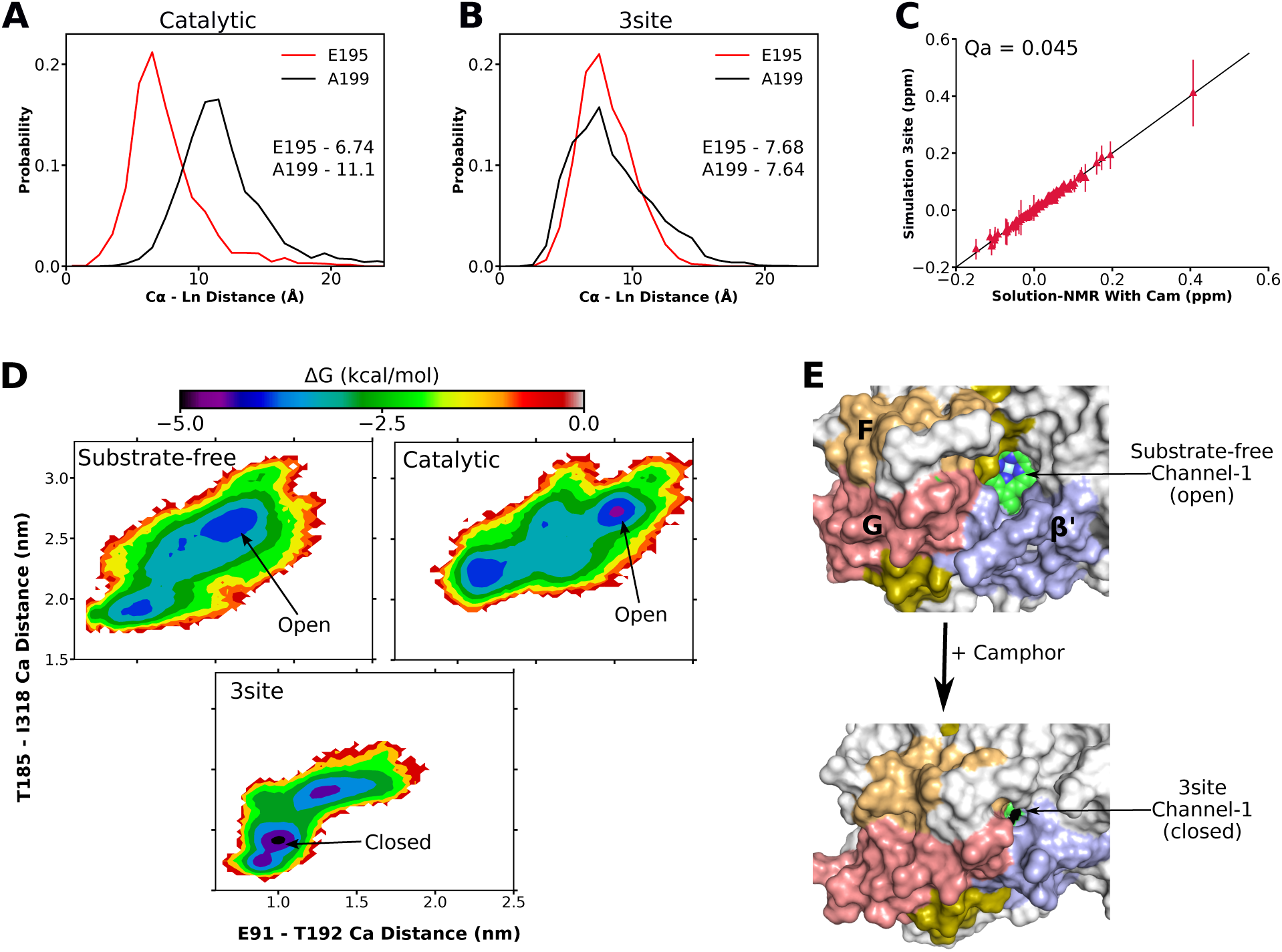
3site state in agreement with known observations: (A,B) The probability distribution of C*α*-Ln distance for residues E195 (red) and A199 (black) obtained by fitting ^1^*H^N^* PCS data (P450cam+Cam) with catalytic (A) and 3site (B) simulation ensembles. Numerical values represents the fitted mean values of C*α*-Ln distance distributions. (C) PCS correlation for observed (solution NMR) and calculated (simulation) PCS for 3site state. Straight line represents perfect correlation. (D) Free energy distribution of conformation of channel-1 for different states of P450cam, depicting open/closed conformations. (E) Open to closed transition of P450cam in 3site state.

Secondly, the channel-1 dynamics as characterised in simulation ensembles were correlated with previous observations. Channel-1 represents an opening near the junction of F, G and *β*′ helices and directly joins the heme center (site of function and catalytic mode binding) to outside bulk solvent. The current and previous studies^9,14,44^ have reported substrate-binding to active site through channel-1. Channel-1 represents the most studied aspect of P450cam and observations from DEER, MD, NMR and crystallographic studies agreed that P450cam explores open conformation of channel-1 in substrate-free state while getting shifted to closed conformation in presence of substrate camphor. ^10,16,29^ The conformation of channel-1 was measured in different states of P450cam, using two distance based metrics i.e., distance between C*α* atoms of T185-I318 and E91-T192 residue pairs (methods). The low to high continuum of these two distances captures the full spectrum of closed to open conformations of channel-1 and hence is being used as the key metric for characterising channel-1 dynamics in the present investigation.

The channel-1 in substrate-free simulation ensemble mostly explored open conformation in agreement with the existing knowledge (Figure 5D, E). Interestingly we found that in presence of substrate camphor, the channel-1 did not transition to closed conformation in catalytic state, instead remained in open conformation (Figure 5D). This also explains why catalytic simulation ensemble did not match with NMR PCS data measured in presence of camphor. However, on the other hand, in the simulation ensemble of 3site state channel-1 significantly shifted towards the closed conformation (Figure 5D,E), in agreement with NMR PCS data. Moreover, binding simulations also captured (Figure 2) the transition of channel-1 from open to closed conformation as the three substrate molecules spontaneously bind to P450cam, (Figure S7A) in real time as a direct consequence of three-substrate bindings, a unique observation not observed in previous binding simulations.^9,14^ The results indicate the simultaneous bindings of three substrate molecules are required for P450cam to transition from open to closed conformation of channel-1. This is in perfect agreement with 3site model that instead of catalytic state, 3site state is the eventual bound state of P450cam.

Further, channel-1 dynamics of present simulation ensembles were remeasured by alternative metrics to corroborate with the previous DEER studies.^10,45^ Channel-1 transitioning to closed conformation in presence of camphor had been measured using DEER approach by spin labelling of S48-S190^10^ or S48-Y179^45^ residue pairs. The distance between C*α* atoms of aforementioned residues pairs was used as closest metric to DEER observations and it was found that channel-1 undergo closing only in 3site state and not in catalytic state, in agreement with 3site model (Figure S7B,C). Overall, we find that only 3site state, in stead of previously known catalytic state, can reconcile and quantitatively interpret the known observations on P450cam’s channel-1, leading to the conclusion that 3site state predominantly exist in solution rather than the catalytic state.

### 3.2 : 3site state establishes substrate-egress mechanism via a different channel

Although 3site state efficiently explained P450cam dynamics, at the first glance it might appear as a state saturated by camphor molecules, with no clear knowledge on the route for product egress. The substrate binding and product-egress paths are generally believed to occur through the same pathway. This apparently presumes that product (catalytic mode) is supposed to egress out through channel-1, which is the substrate binding pathway in the active site. Additionally, previous studies targeting product egress paths in P450cam reported channel-1 as either sole or major product egress pathway.^46–48^ But in 3site state, the waiting mode sits on top of catalytic mode within channel-1 and hence it appears impossible for catalytic mode to egress out through channel-1 (Figure 6A). Recently as an alternative, mechanism with separate binding and egress pathways (channel-1 and channel-2 respectively, Figure 6A) had also been hypothesized for P450cam to account for its very fast kinetics. ^14^ For an unbiased understanding of the product egress route in P450cam, both the channel-1 and channel-2 were assessed for their viability as potential egress routes for substrate bound in catalytic mode. As derived by spectroscopy based measurements, P450cam has previously been observed to exhibit remarkably fast kinetics with a *k_off_* of 15 *s^−^*^1^ corresponding to 67 ms of substrate residence time in active site.^30^ The unbinding simulations of catalytic camphor were carried out through different channels using infrequent metadynamics approach and unbiased residence time was calculated and used to confirm the fast kinetics of P450cam.

**Figure 6:**
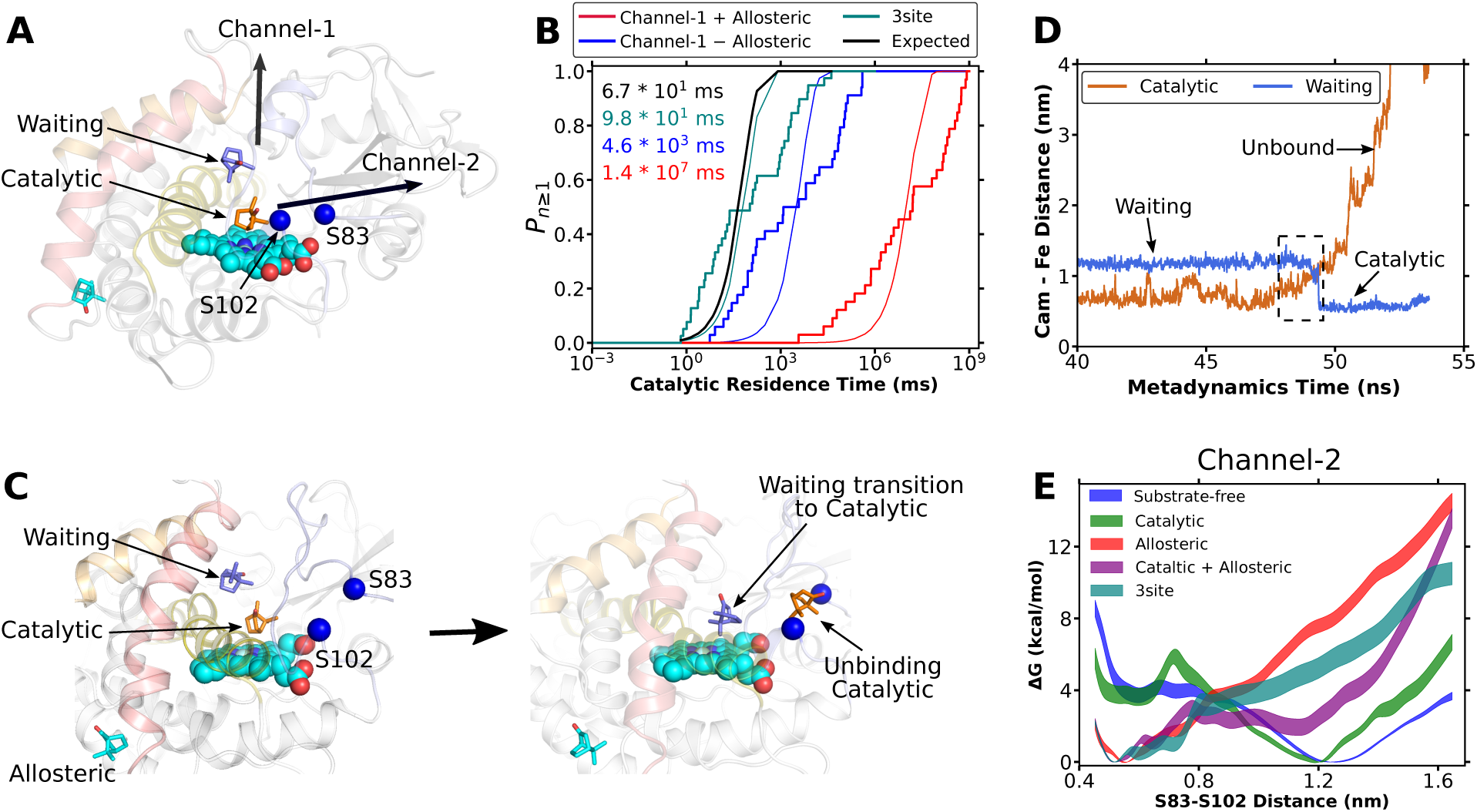
Product egress in 3site state: (A) Channel-1 and channel-2 as potential egress routes of catalytic mode. (B) Probability of unbinding of catalytic mode through different channels of P450cam. Numerical values represents the residence times of catalytic mode. (C,D) Mechanism of product egress in 3site state showing ‘waiting mode transitioning to catalytic mode’ in concert with catalytic mode unbinding. (E) Underlying thermodynamics of channel-2 opening in different states of P450cam. S83-S102 distance ≤ 0.8 nm represent the closed conformation, while larger represents the open conformation. Width of curve represents the error.

Firstly to assess the feasibility of the channel-1, which happens to be the entry-path of the substrate, as the egress route, the waiting mode was not considered and unbinding simulations of catalytic mode were performed with and without the allosteric mode. The distance between camphor (catalytic) and Heme-Fe atom was used as collective variable such that the unbinding shall occur only through channel-1. 50 such unbinding simulation replicates were used to calculate the residence time of catalytic mode as a measure of time taken by camphor to egress out through channel-1. The residence time of catalytic mode through channel-1 in presence of allosteric mode was found to be 1.4 ± 0.2 × 10^7^ ms (Figure 6B, S8A, Table-1). Additionally, the residence time of catalytic mode was also calculated without allosteric mode which comes out to be 4.6 ± 1.0 × 10^3^ ms (Figure 6B, Table-1). Clearly, even if the waiting mode was not considered, unbinding of substrate at catalytic mode through channel-1 is very slow as it required significantly large time to egress out (as compared to expected 67 ms). Hence, we concluded that channel-1 as an egress route is not in agreement with very fast kinetics observed in P450cam and hence most likely is not the potential product egress route.

For an unbiased understanding of product egress route, we devised a new collective variable to simulate unbinding in the 3site state. The distance between camphor (catalytic mode) and catalytic center of P450cam (Heme-Fe, D251, T251, V396, methods) was used as collective variable such that unbinding can occur through both channel-1 or channel-2. The presence of waiting mode in 3site state shall restrict unbinding through channel-1 and hence channel-2 might be preferred. Total 50 unbinding replicates were performed, in 43 replicates the catalytic camphor egressed out through channel-2, while in remaining 7 replicates the catalytic camphor chose channel-1 as egress route. The overall residence time of catalytic camphor in 3site state was found to be 98 ± 7 ms, in very close agreement with expected 67 ms (Figure 6B, Table-1). The closer look of unbinding mechanism in 3site state revealed interesting features. Starting with the 3site state, the waiting mode waits for its turn by remaining at 1.19 nm away from heme-Fe (Figure 6C). As soon as the substrate at the catalytic mode unbinds through channel-2, the substrate at waiting mode transitions to the position vacated by the original substrate in catalytic mode. (Figure 6C,D). This mechanism of “waiting mode transitioning to catalytic mode” upon product egress explains the fast kinetics observed in P450cam, which can be reconciled only in 3site state. Contrastingly, unbinding through channel-1 resulted in simultaneous unbinding of waiting mode in few cases, thereby defeating the purpose of 3site state (Figure S8B). Hence, we concluded that channel-2 is the major product egress route in P450cam, while channel-1 might serve as a minor product egress route.

Initially postulated by Follmer et al, ^14^ the channel-2 flanked by flexible loops on both sides (marked by S83 and S102) is a roughly orthogonal route with respect to channel-1, thereby allowing safe egress route to catalytic mode without interfering with waiting mode (Figure 6A). To investigate the closing/opening equilibrium of channel-2 in different states of P450cam, the underlying free energy change for channel-2 opening was calculated as a function of the distance between S83 and S102 Ca atoms. A S83-S102 distance less than 0.8 nm was referred as closed conformation of channel-2 and that greater than 1 nm as its open conformation. The channel-2 opening was found to be free-energetically favourable in case of substrate-free and substrate-bound catalytic state (Figure 6E). Upon addition of substrate in the allosteric and/or waiting modes, the channel-2 opening becomes energetically unfavourable such that opening of channel-2 in 3site state shall required 3.88 ± 0.55 kcal/mol energy (Figure 6E), in direct disagreement with observations of Follmer et al.^14^ which are potential artifacts of hydrogen mass repartitioning (HMR) approach^49^ that was used for simulation. (detailed comparison with HMR in SR-2) The three binding modes controlled the I helix movements, which in turn controlled the channel-2 opening, hence leading to different energetics of channel-2 in different states of P450cam (detailed allosteric mechanism in SR-2). Overall, in the final 3site state, the channel-2 remain in closed conformation and product egress shall require its opening as observed in unbinding simulations (Figure S8C).

### 3.3 : 3site state possessed regio-selective conformations of substrate camphor

Besides camphor, P450cam also binds another small substrate i.e., *O*_2_ molecule at heme-Fe, followed by final hydroxylation reaction. P450cam is highly regio-selective as revealed by spectroscopic analysis of camphor products, such that the hydroxylation occurs specifically only on C5 carbon of substrate camphor ^39,50^ (Figure 7A). Since 3site is hereby claimed to be the final bound state for P450cam, then this state in presence of *O*_2_ molecule should have regio-selective conformations of catalytic binding mode (the substrate that will undergo reaction). To understand whether the regio-selective nature of P450cam can be explained in catalytic or 3site state, the conformational preference of substrate camphor (catalytic binding mode) was measured for both the state in absence/presence of *O*_2_ molecule. In order to account for the binding of *O*_2_ molecule, the separate simulation ensembles of catalytic and 3site states were generated with oxy-heme (Heme-Fe atom coordinated to *O*_2_ molecule, methods).

**Figure 7:**
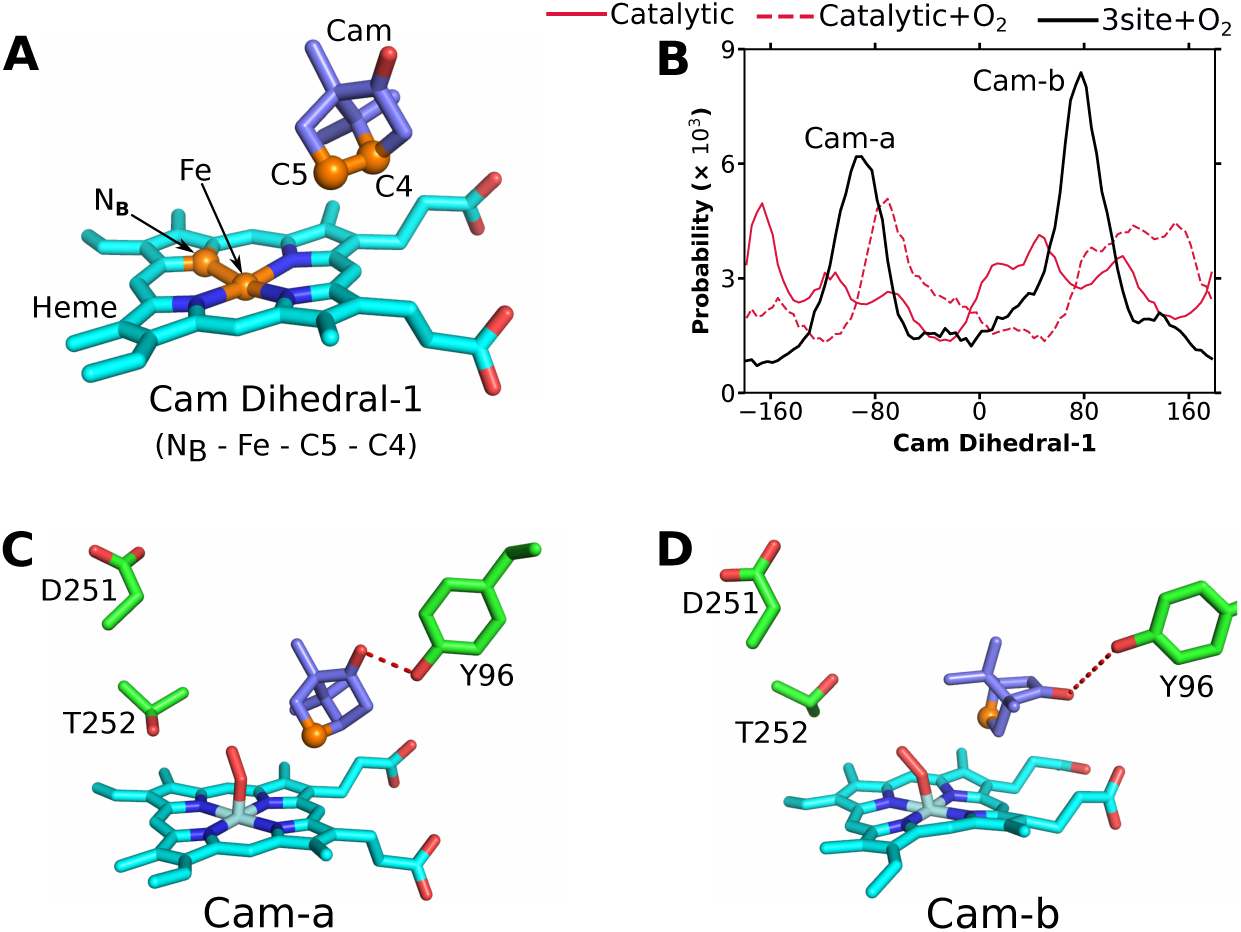
Regioselectivity in 3site state: (A) The catalytic mode above heme moiety depicting C5 atom and cam Dihedral-1 used to quantify conformational preference of substrate camphor. (B) Probability distribution of cam Dihedral-1 in different states of P450cam. (C,D) The representative snapshots corresponding to regio-selective *cam-a* and *cam-b* conformations of catalytic mode.

Camphor hydroxylation occurs on C5 carbon atom of camphor, which should be positioned towards catalytic center, hence a dihedral (Cam Dihedral-1, *N_B_*-Fe-C5-C4) was designed to quantify the orientation of C5 relative to catalytic center (Figure 7A). As measured by Cam Dihedral-1, the substrate camphor in catalytic simulation ensemble does not exhibit significant preference for any particular conformation within the active site (Figure 7B). Upon addition of *O*_2_ molecule in catalytic state, substrate camphor exhibit relatively better but broad preference for conformations corresponding to Cam Dihedral-1 values of around -80 and 80-160*°* (Figure 7B). Interestingly, substrate camphor in 3site+*O*_2_ ensemble markedly exhibit conformational preference for two distinct poses with Cam Dihedral-1 values of around -85 and 80*°*, corresponding to two distinct conformations namely *cam-a* and *cam-b* (Figure 7B-D). Both conformations *cam-a* and *cam-b* revealed two important features; first, O atom of camphor found to orient toward Y96 sidechain and hence able to formed hydrogen bond with it; second, the C5 atom of camphor was found to orient toward catalytic center and hence in favourable conformation to undergo hydroxylation reaction (Figure 7C,D). Results indicate that 3site state possessed regio-selective conformations of substrate camphor (catalytic mode) and hence is the functional bound state of P450cam that undergoes reaction (also see SR-4); while catalytic state did not. Further, although small, *O*_2_ was found to play an important role in substrate regio-selectivity as even the 3site state without *O*_2_ did not posses regioselective preference (Figure S9A). Additionally, carbon atoms C7 and C9 were known to play important role in regio-selectivity as another substrate norcamphor (missing C7 and C9) exhibits decreased regioselectivity.^39,50^ Hence we devised another dihedral (Cam Dihedral-2, *N_D_*-Fe-C7-C9, as used previously^51^) to quantify the conformational preference from the perspective of C7 and C9 carbon atoms (Figure S9B). For Dihedral-2 also, only 3site+*O*_2_ state exhibit regio-selective conformational preference (Figure S9C).

### 3.4 : Redox Partner Putidaredoxin retains closed conformation of 3site state

For successful reaction, P450cam further requires 2*e^−^* from redox partner protein putidaredoxin (Pdx). It had been claimed by crystallography^23^ and MD^27^ based observations that Pdx induced back P450cam+Cam to open conformation of channel-1 where final reaction occurs in open environment. Contrastingly, NMR^29^ based observations claimed that Pdx maintained the P450cam+Cam in closed conformation and reaction occurs in isolated closed environment. The contrasting observations led to controversial role of pdx whether it can open channel-1 or not. To evaluate the potential effector role of Pdx in light of 3site model, the additional simulation ensembles of various states of P450cam bound to Pdx were generated (methods) and conformation of channel-1 in different states of Pdx bound P450cam were measured. It was found that in substrate-free state, P450cam remained largely in open conformation both with or without Pdx (Figure 8A, S10). For 3site state, which transition to closed conformation without Pdx, P450cam+Pdx also transition to closed conformation (Figure 8A, S10). The one-to-one comparison of channel-1 in different states of P450cam with and without Pdx revealed that Pdx did not stabilize the open conformation of channel-1 in any of the substrate-bound state of P450cam, instead induced closed conformation in some state of P450cam (Figure S10). Together, the P450cam + substrate + Pdx i.e., the substrate and Pdx bound P450cam (3site+Pdx) in closed conformation of channel-1 indicates that the Pdx bound P450cam has channel-1 in closed conformation (Figure 8B), in agreement with the previous NMR-based observations that reaction occurs in isolated closed environment,^29^ while in disagreement with crystal and MD based observations.^14,23^

**Figure 8:**
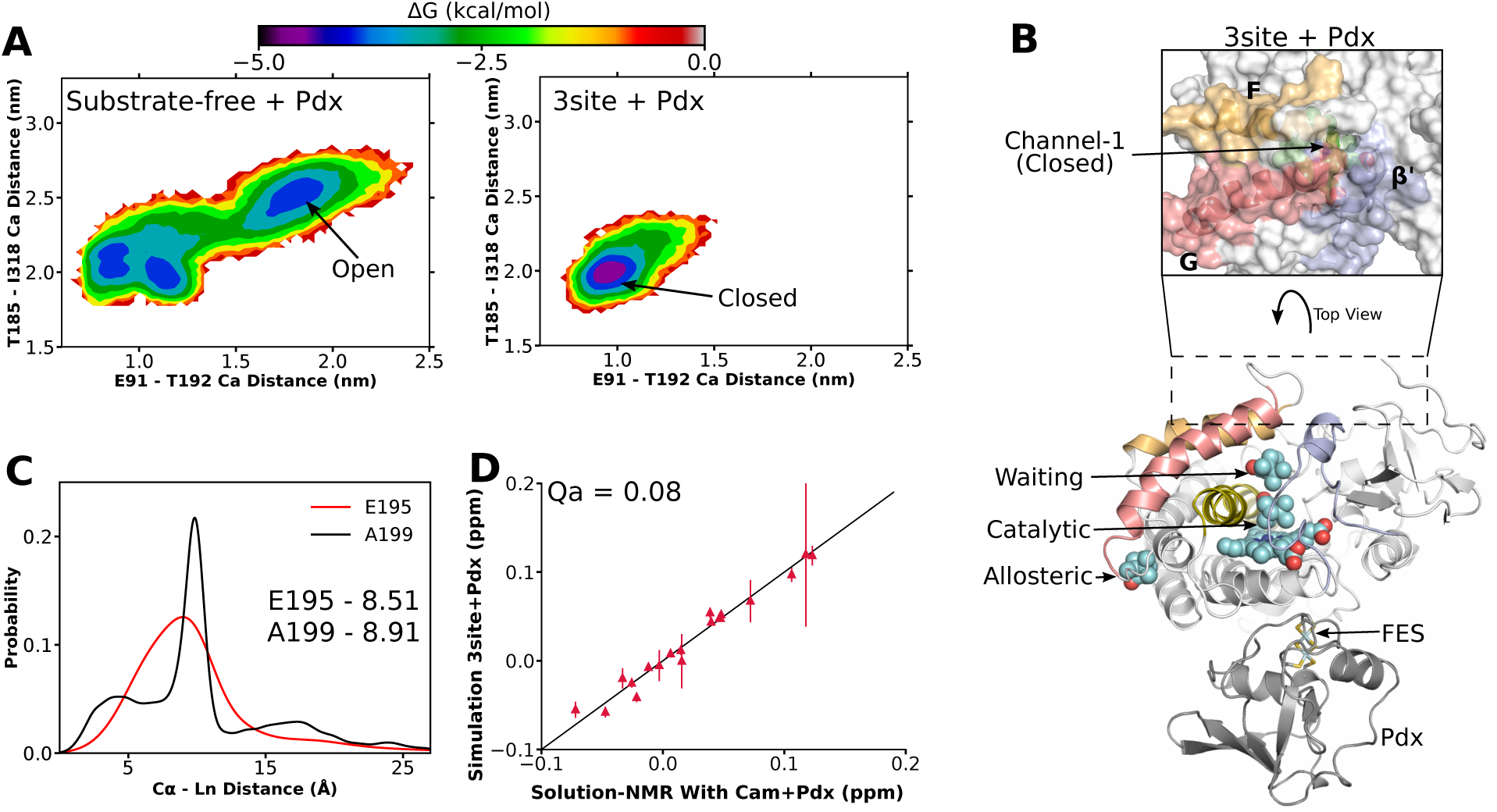
Pdx and substrate bound P450cam is closed: (A) Free energy distribution of channel-1 depicting open/closed conformation in different states of Pdx bound P450cam. (B) Closed conformation of channel-1 in 3site+Pdx state. (C) The probability distribution of C*α*-Ln distance obtained by fitting ^1^*H^N^* Leu PCS of P450cam (in presence of both substrate and Pdx) with 3site+Pdx simulation ensemble. Numerical values represents the fitted mean values of C*α*-Ln distance distributions. (D) PCS correlation for observed (solution NMR) and calculated (3site+Pdx simulation ensemble).

Additionally, attempts have been made to correlate the contrasting observations from crystallography or MD with that of solution NMR. Unfortunately till date, no Pdx+substrate bound P450cam structure either from crystallography or MD simulations has been able to match the solution NMR PCS data^29,42^ potentially because only one binding mode (catalytic) was considered. Hence, we checked whether the novel 3site+Pdx state can match the NMR PCS data. Therefore, the previously reported ^1^*H^N^* Leu PCS of P450cam measured in presence of substrate and Pdx were taken. ^29^ The P450cam structures after every 100 ps time interval from 3site+Pdx simulation ensemble were used to calculate the simulation PCS (methods). The simulation and NMR PCS fitted perfectly with C*α*-Ln distance of 8.51 and 8.91 °*A* for E195 and A199 respectively, i.e., within the expected range of 7-9 °*A* (Figure 8C). The simulation PCS were also found to be in perfect agreement with NMR PCS (Figure 8D). The difference between the simulation and NMR PCS (Qa) was found to be significantly low with a value of 0.08. Overall, the results conclude that Pdx does not stabilize the open conformation of P450cam and P450cam+Pdx transitions to closed conformation upon binding of substrate camphor at catalytic, waiting and allosteric binding modes (3site state).

### 3.5: 3site state can be modulated by L166A and T217V mutations

In the end, to check the potential modulations of 3site state, the site specific mutants were designed. Apart from known important residues in P450cam, this work suggest two additional residues which were found important for P450cam’s functioning; firstly, residue L166 which is part of both allosteric site and E-I cavity and play important role in allosteric coupling of active and allosteric sites (SR-3, Results-2.1, also previously observed in NMR chemical shifts^52^), and secondly, residue T217 which formed hydrogen bond with allosteric mode and stabilize it (Results-1, Figure-1, S2). To further confirm the importance of L166 and T217 residues, in-silico site specific mutants were created. For both mutants, residence time of allosteric mode was measured and compared with wildtype observation (Table-1).

Firstly, residue L166 if mutated to smaller alanine (L166A) stabilized the *I^th^*-kink and reduced the allosteric pocket volume to 82.3 °*A*^3^ as compared to 234.8 °*A*^3^ in wildtype (SR-3). In simulations of allosteric and catalytic+allosteric states of L166A, the allosteric camphor spontaneously left the reduced allosteric site within very short simulation time. The spontaneous unbinding times of allosteric mode observed in 56 simulations were used to calculate the residence time of allosteric mode in L166A mutant. It was found that allosteric mode was unable to reside in allosteric site and exhibit the residence time of only 0.166 ± 0.003 *μ*s i.e., 85 times decrease compared to wildtype (Figure 9C, Table-1). Overall, L166A mutant rendered P450cam’s allosteric site incapable of binding (detailed in SR-3).

**Figure 9:**
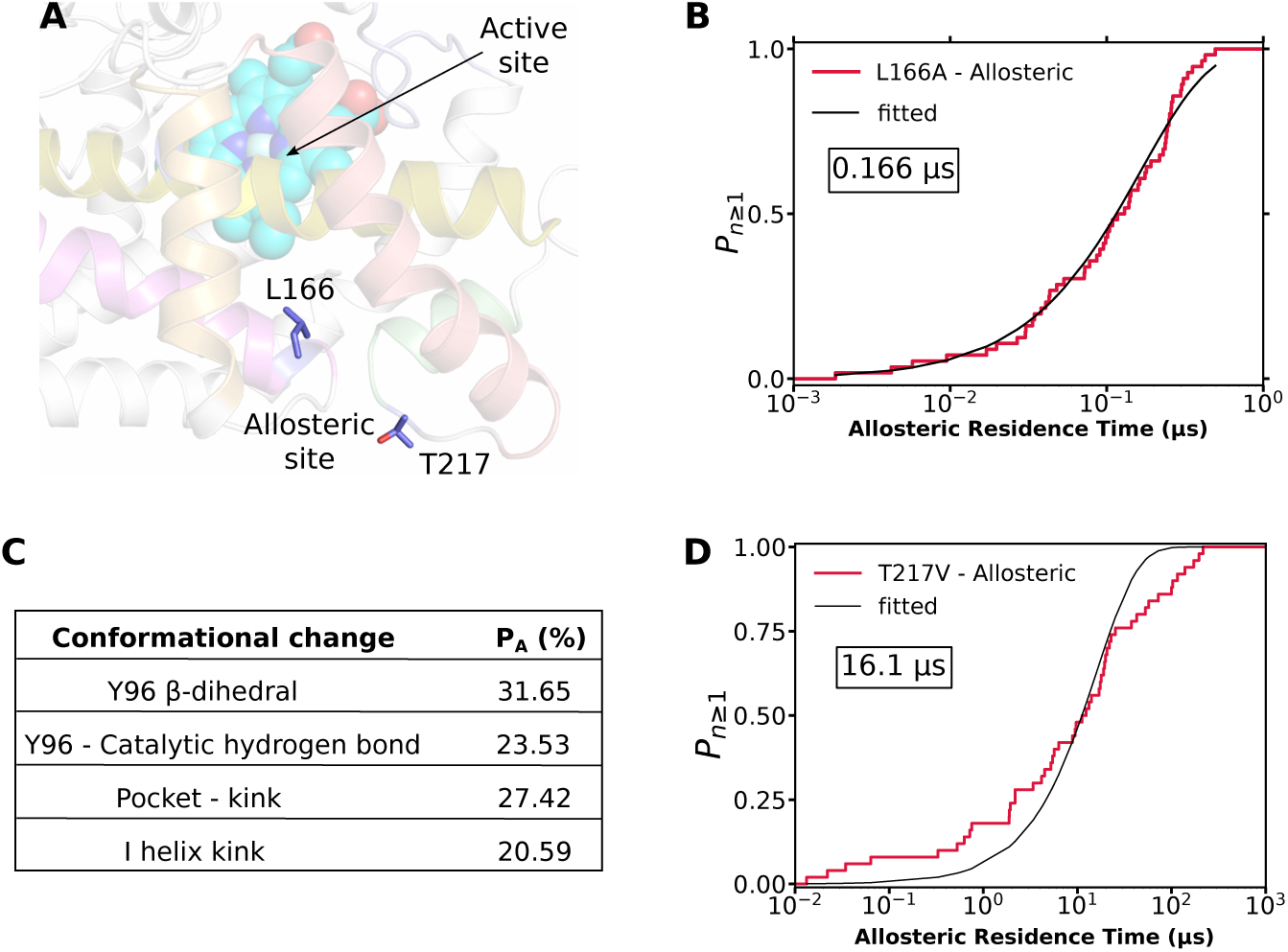
Modulations of 3site state: (A) Residues L166 and T217 in allosteric site. (B) Probability of allosteric mode unbinding in L166A mutant. Numerical value represent the residence time of allosteric mode. (C) Percent contribution of allosteric mode to functionally important conformational changes within the active site. *P_A_* represents the percent of total conformational change that is induced by allosteric mode. (D) Probability of allosteric mode unbinding in T217V mutant.

Additionally, as measured by NMR chemical shifts, L166 was previously known to be one of the most perturbed residues of P450cam upon addition of camphor, but without the knowledge of allosteric site or 3site state. ^52^ Further, as measured by NADH usage, the L166A mutation decreased the function (at active site) by 30%.^52^ In light of 3site state, this work suggests that L166A mutation rendered allosteric site unable to be occupied by substrate. Since the three binding modes function synergistically, 3site model implied that the previously observed decreased function in L166A mutant should come from allosteric mode which shall be equal to 30% (the observed decrease in function in L166A^52^). As the three binding modes additively contributes to function in the active site (Figure 4), the separate contribution of allosteric mode was calculated (*P_A_*). For any functionally important conformational change in active site, *P_A_* represent the portion of conformational transition induced by the allosteric mode. For instance, the contribution of allosteric mode in transitioning Y96 (in active site) from *conf-a* to *conf-b* conformation was calculated (methods). Interestingly, it was found that allosteric mode roughly contributed 30% of various functional conformational changes within the active site (Figure 9D). For instance, allosteric mode contributes 31.65% of the total transitioning of Y96 conformation from *conf-a* to *conf-b*. Approximately 30% of total conformational changes contributed by allosteric mode shall not be present in L166A mutant, inturn decreasing the function in active site. Overall, the previously observed decreased function of distant L166A mutation can be easily explained by 3site model.

Secondly T217, which formed hydrogen bond with allosteric mode (Figure 1, S1), was mutated to non-hydrogen bonding valine to confirm the role of Cam-T217 hydrogen bond in binding of allosteric mode. The simulation ensemble of allosteric state with T217V mutation was generated and 50 metadynamics simulations were performed on allosteric mode. The T217V mutant did not destablized the allosteric mode which remain bound to allosteric site during the unbiased simulations. Contrasting to the expectation, the allosteric mode was found to be more stable in T217V mutant giving the residence time of 16.1 ± 0.7 *μ*s as compared to 4.1 ± 0.2 *μ*s in wild type (Figure 9C, S11, Table-1). This indicates that Cam-T217 hydrogen bond is not crucial for binding at the allosteric site, instead it might be the hydrophobic nature of allosteric site that is crucial for binding. This shall be in agreement that the substrates of P450cam and other P450s all have large hydrophobic moieties. ^53^

## Discussion

This work reports a discovery of novel state in P450cam where three substrate camphor molecules simultaneously bind to P450cam resulting in 3site state. For the first time, a bound state involving three interlinked binding modes is proposed for cytochrome P450 family, which is clearly distinct from previously proposed bound states involving one or two binding modes. Interestingly enough, the three binding modes are allosterically linked with each other and function synergistically such that three simultaneous bindings are crucial to fucntion. The 3site state is consistent with the plethora of precedent experimental measurement including NMR, crystallography, DEER, product egress kinetics, spectroscopy and mutagenesis. Most importantly, it was found that 3site state shall predominantly exist in solution as the final bound state of P450cam. Based on the insights gathered, the overall functioning of P450cam is summarized in Figure 10: starting with the open conformation of channel-1 in substrate-free state, three camphor molecules simultaneously bind to P450cam yielding the closed 3site state. Although the 3site state although closed conformation of channel-1, D251 residue remained free of its salt bridges with R186 and K178, hence retaining functionality (SR-4). Upon regioselective hydroxylation of catalytic mode, product egress occurred through channel-2 with simultaneous shifting of waiting mode to catalytic mode and another substrate binds to waiting mode. Meanwhile, the binding/unbinding at allosteric site continues simultaneously with events in active site. Additionally, Pdx binds P450cam either before/during/after camphor binding and donate electron without opening channel-1.

**Figure 10:**
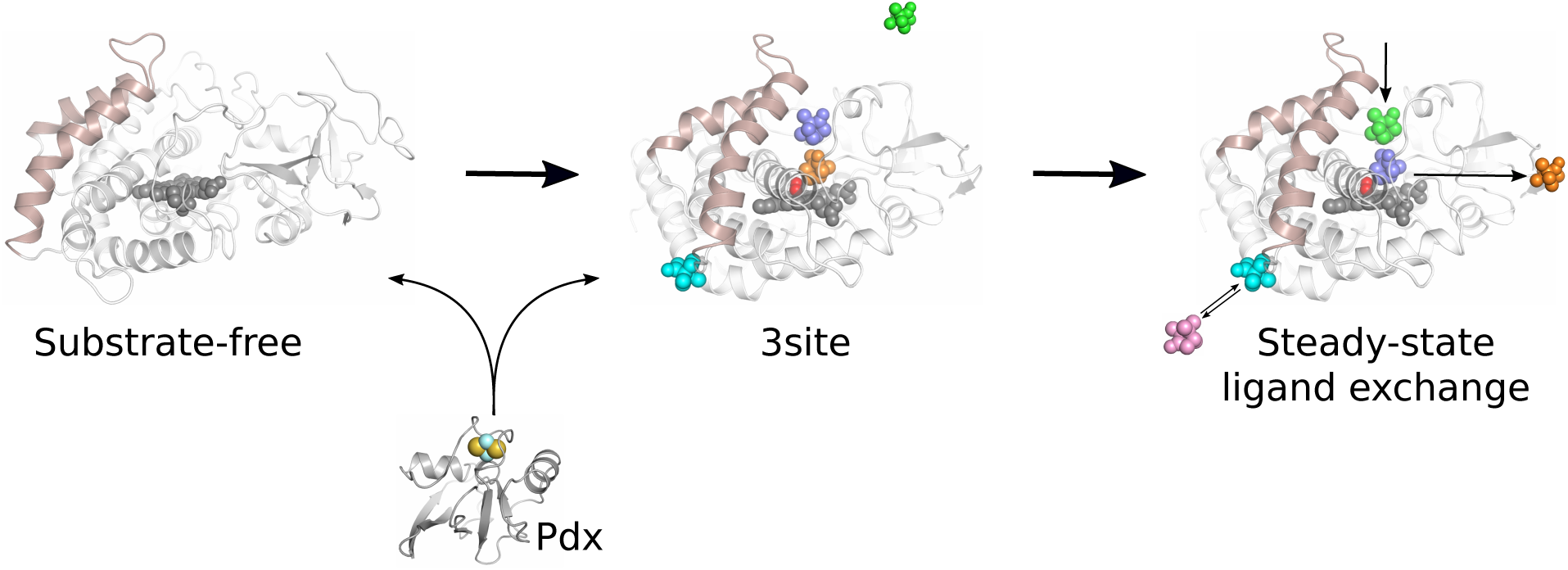
Proposed functioning mechanism of P450cam: P450cam in substrate-free state with open conformation of channel-1 binds 3 substrate molecules at catalytic (orange), waiting (purple) and allosteric (cyan) modes, yeilding 3site state in closed conformation. The 3site state is functional and allows regioselective hydroxylation of substrate in active site (catalytic mode). The product (catalytic mode) egress out through channel-2, vacating catalytic mode which then occupied by waiting mode while another substrate (green) from solution binds at the waiting mode. Simultaneously, binding and unbinding goes on at the allosteric site. Upon unbinding, another substrate (violet) binds to allosteric site. Pdx also bind to P450cam and donate electron for reaction in active site without opening channel-1.

Via reconciling 3site state with previous P450cam’s observations, this work proposes that there is a potential chance that it is the 3site state which predominantly exist in the solution and the previous on 3site ensemble rather than the catalytic state. This was justified by explicitly calculating the various observables for catalytic and 3site states, where 3site state demystified all of them, while catalytic state did not. But this intrigued a conundrum, specifically why only the catalytic, and not allosteric and waiting, modes were observed in crystal structures so far. Gratifyingly, existence of additional binding modes (other than catalytic) for P450cam has been postulated before, but not known yet. Additionally, the waiting mode has been observed in one crystal structure (4JX1),^23^ but was not recognized as separate binding mode as the structure garner controversy for Pdx’s role. Similarly for allosteric mode, the unusual nature of a region at base of F/G helices (allosteric site) has been observed before in NMR experiments.^29^ Additionally, a large unknown electron density (not water) at allosteric site has been observed in one crystal structure (5IK1).^26^ Overall, despite multitudes of postulation and indirect evidences of multiple binding modes, such a structure has not been determined yet. The observation of waiting state at large shall depend on allosteric mode, since allosteric mode is required for stable waiting mode. The allosteric mode might not be trapped in crystals owing to its low binding energy or residence time and fast dynamics. Crystals are well known for their missing regions such that portions of structures having high flexibility (beta factor) cannot be determined. The allosteric mode is distinct in itself because it has low binding energy of -2.66 kcal/mol and very high binding/unbinding cycle owing to 0.14 and 14.1 *μ*s of binding-unbinding times respectively. Crystal structure for such a mode can be determined by obtaining crystals at high concentration. This is in agreement that most crystals were obtained at 1mM concentration of camphor, while crystal obtained at 2mM concentration (5IK1) exhibit unusual density at allosteric site. ^14,26^ Therefore, we propose that recrystalizing 4JX1 type crystal at higher concentration might yield the till now hidden 3site state.

The archetypal P450cam had served as primary model for ubiquitous cytochrome P450s owing to largely conserved sequence, P450-fold and heme chemistry in CYP superfamily. ^1^ The 3site state observed in P450cam (CYP101A1) functions through a crucial kink in I helix, which allows the allosteric linkage between the active and allosteric sites. The kink in I helix generates due to presence of low helical propensity glycine residues at 248 and 249 positions, just before the functionally important D251 and T252 (SR-1-3). The sequence analysis reveals striking conservation of glycine residue at two residues preceding to functional D251 in all the P450s ranging from prokaryotes to humans (Figure S11). This is in agreement that crystal structures from different kingdoms were determined with *I^th^*-kink and allosteric pocket is also conserved in many P450s.^15^ The absolutely conserved glycine, even more than functional D251, clearly indicates that the *I^th^*-kink might be functioning towards allosterically linking the active and allosteric sites in all cytochrome P450s. Gratifyingly, multiple binding modes has been postulated for other P450s and a ‘waiting like binding mode’ has been determined in at least one crystal structure (PDB id - 1W0F).^17–22^ Specifically, the non-Michaelis behaviour of many P450s had been known for atleast two decades,^21^ but the molecular mechanism involving postulated multi-binding mode state is not known yet. This is compelling that hereby reported 3site state might be functioning in other P450s also. Additionally, the 3site state with three different yet interlinked binding modes can be the fascinating system for allosteric studies.

## Methods

### Unbiased binding simulations

The crystallographic structure of 4JX1^23^ in open conformation of channel-1 was used as starting structure. One copy of P450cam corresponding to chain A of 4JX1 (residue 10-414) with heme was used. Protonation states of all amino acids corresponds to neutral pH, except D297. D297 was buried in active site and formed hydrogen bond with one of the heme propionate moieties, hence modelled with neutral (protonated) side chain. To perform the unbiased binding simulations, the substrate-free open P450cam was placed at center of cubic simulation box with 10 °*A* minimum distance between protein surface and box. The system was solvated with around 22500 water molecules and *Na*^+^ ions were added to overall neutralize the system. Four substrate camphor molecules were randomly placed in bulk solvent media corresponding to camphor concentration of 8.8 mM and experimentally suggested 4:1 substrate:protein ratio. ^54^ Overall, the total system included around 74000 atoms (Figure S12). The P450cam protein, heme and ions were parametrized using all-atom CHARMM36 forcefield with the backbone CMAP correction. ^55^ For substrate camphor, the CHARMM compatible parameters derived using GAAMP^56^ from our previous work were used.^9^ Water molecules were modelled using charmm-TIP3P water model.^57^ All MD simulations were performed using GPU coupled gromacs 20xx packages. ^58^ The simulations were performed at an average temperature of 303.15 K, maintained using Nose-Hoover thermostat with a relaxation time of 1.0 ps.^59,60^ Long range electrostatistics and Lenard-Jones interactions within 1.2 nm cut-off were calculated using PME summation ^61^ and Verlet cutoff scheme^62^ respectively. Hydrogen bonds were constrained using LINCS algorithm^63^ and water hydrogen bonds were fixed using SETTLE algorithm.^64^ Before production run, system was energy minimised and equilibrated for 5 ns in NVT and 5ns in NPT ensemble using position restraints. In the end, the restraint free production run was performed in 12 replicates. The substrate camphor molecules were allowed to unbiasedly sample the P450cam conformational space and search its stable binding modes for a maximum of 4 *μ*s. The binding process at any instant was checked by two metrics: the center of mass distance between camphor and Fe atom of heme moiety was used to detect the substrate location in active site; while the minimum distance between heavy atoms of camphor and L166 was used to detect the substrate location nearby allosteric site (a non-native site, see results). The simulations were stopped once stable binding(s) were achieved for 100’s of nanoseconds. Additionally, the unbiased binding simulations reported in our previous work were also used in this work.^9^ Briefly, the binding simulations were started with closed conformation of P450cam (corresponding to PDB ID 2CPP^5^), placed at center of dodecahedron box and filled with solvent, *Na*^+^ and 4 substrate camphor molecules. 3 independent binding simulations were performed, similar to above with only difference that starting configuration of P450cam was in closed conformation. Detailed description was provided elsewhere. ^9^

### Simulation ensembles of different states of P450cam

18 different states of P450cam were defined based on hereby described 3site model, presence/absence of Pdx, heme-*O*_2_ co-ordination or mutations (Figure 2, S13). For each state, a large simulation ensemble was generated. To generate a simulation ensemble, the starting configuration with either desired binding mode(s), Pdx, heme-*O*_2_ or mutation was taken in center of cubic box and filled with water and *Na*^+^ ions. Parameterization was performed similar to binding simulations. For iron-sulfur cluster in Pdx, the ab-initio generated forcefield parameters of *Fe*_2_*S*_2_ and coordinated cysteines were taken. (Figure S14). Three independent simulations to a maximum of 1 *μ*s were performed, unless mentioned otherwise. All the independent simulations (after 0.35 *μ*s) were combined to generate the final simulation ensemble of each state (See supplemenatry methods (SM 1-2)).

### Calculation of NMR pseudo contact shifts from MD simulation ensembles

To verify that structures of in-silico simulation ensembles do exist in in-vitro solution state, the Pseudo contact shifts (PCS)^65^ measured in solution NMR were correlated with predicted PCS derived from simulation ensembles. Four sets of TROSY-HSQC derived PCS were used in this work, from two previously reported studies.^24,29^ The four amide PCS sets were; (i) ^1^*H^N^* Leu PCS of substrate-free P450cam, (ii) and (iii) ^1^*H^N^* Leu PCS and ^1^*H^N^* PCS of P450cam in presence of substrate camphor, and (iv) ^1^*H^N^* Leu PCS of P450cam in presence of both substrate camphor and redox partner Pdx. The detailed methods were provided elsewhere.^24,29^ Importantly, the E195C/A199C/C344A mutant of P450cam was labelled with either paramagnetic (Yb) or diamagnetic (Lu) tag through CLaNP-7 attached to C195 and C199 of G helix. The ^1^*H^N^* PCS were calculated as difference between chemical shifts of paramagnetically and diamagnetically tagged samples.

From simulation ensembles of P450cam with or without substrate camphor, configurations after every 1 ns were taken, while for simulation ensembles involving Pdx, configurations after every 100 ps were taken for predicting PCS. The Δ*χ* tensor for each of the simulation ensembles was determined using ensemble average fit, where single tensor was simultaneously fitted to all frames of the simulation ensemble. The fitting was performed using paramagpy library in python.^66^ The position of paramagnetic tag was determined using singular value decomposition (SVD) grid search, within a sphere of 10 °*A* radius having 10 fitting points per radius, by using the observed tensor (solution NMR) as the starting point. Subsequently, Δ*χ* tensor for any simulation ensemble was determined using the corresponding set of observed PCS data and the PCS for the simulation ensembles were calculated using fitted Δ*χ* tensor. The reliability of Δ*χ* tensor was analyzed according to ensemble average distance between C*α* atoms of E195, A199 (attachment site of CLaNP-7) and predicted position of paramagnetic tag (Ln), i.e., C*α*-Ln distance. The probability distribution of C*α*-Ln was estimated using kernel density estimation^67^ with bandwidth selection using Scott’s rule of thumb^68^ (the mean remain unchanged from raw distances). The C*α*-Ln distance was expected to be 7-9 °*A* due to steric constraints as paramagnetic tag was attached to protein residues through CLaNP-7 molecule.^69^ Fitted tensors having mean C*α*-Ln distance within 7-9 °*A* range were considered reliable and analysed further. Finally, the fitting between observed PCS from solution NMR (obs) and calculated PCS from simulation ensembles (calc) was measured using Q score^43^ as per eq(1):

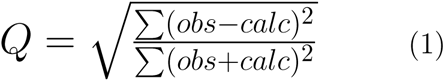

### Free energy calculations

Firstly, binding free energy of catalytic binding mode in presence/absence of allosteric mode was calculated using umbrella sampling approach. The binding energy was calculted as a function of camphor-hemeFe distance, ranging between 0.45-2.00 nm and discretized into 32 umbrellas. Each umbrella was sampled for 25 ns, reweighted using WHAM approach and error analyzed using bootstrapping. Binding energy was calculated as difference between bound (minima of curve) and unbound (2.00 nm). Secondly, thermodynamics of channel-2 opening/closing transition in 5 different states of P450cam (substrate-free, catalytic, allosteric, catalytic+allosteric and 3site) was mapped by umbrella sampling approach. The opening of channel-2 was measured by distance between Ca atoms of S83 and S102, ranging from 0.45-1.65 nm and discretized into 25 umbrellas. Each umbrella was sampled for 25 ns and analyzed as mentioned above. Thirdly, binding energy of allosteric mode was measured by natural log of probability distribution of camphor locations in binding simulations detected as a function of minimum distance between heavy atoms of camphor and L166 (detailed in SM-3).

### Calculation of substrate residence time

Well tempered Infrequent Metadynamics^37,70,71^ (detailed in SM-4) approach was used to calculate the residence time of camphor in different binding modes. For catalytic binding mode, the residence time was calculated in 3 states, i.e., in absence of other modes, in presence of allosteric and both allosteric and waiting modes. For waiting mode, residence time was calculated in 2 states, i.e., in presence/absence of allosteric mode. For allosteric mode, residence time was calculated in 5 states, i.e., in absence of other binding modes, in presence of catalytic mode, in presence of catalytic and waiting modes and with L166A and T217V mutations. For L166A-allosteric, spontaneous unbindings were used. For each case, 50 unbinding simulations were performed (56 for L166A-allosteric). The empirical cumulative distribution function of transition times was cumulative distribution function of poisson distribution^38^ and residence time was calculated by curve fitting and goodness was measured by Kolmogorov-Smirnov test.^72^

Additionally, 3D architecture, volume or constituent residues of active and allosteric sites were analyzed using Mdpocket algorithm and in-house Python codes (SM-5). E91-T192, T185-I318, S48-S190 and S48-S179 C*α* distances were defined as order parameters for channel-1 as detailed in SM-6. Contribution of allosteric mode to function in active site was calculated as difference in conformational change in catalytic+allosteric and catalytic states, as detailed in SM-7.

## Supporting information

supplemental figures, table and extended results

## Supplemental Information

Details of method and supplemental figures and extended results.

## Acknowledgements

We thank Dr. Marcellus Ubbink and Simon Skinner for generously sharing useful pseudo contact shift data obtained from their NMR measurements on P450cam. We thank Dr. Talis Uelisson da Silva for useful discussion regarding parameter generation of iron-sulfur cluster. We also thank Dr. Bibhab Majumder and Dr. Bhupendra Dandekar for their useful discussions. This work was supported by shared computing resources obtained from TIFR Hyderabad, India. We acknowledge support of the Department of Atomic Energy, Government of India, under Project Identification No. RTI 4007. JM acknowledges Core Research grants provided by the Department of Science and Technology (DST) of India (CRG/2019/001219).

## Notes

### Competing Interest Statement

The authors have declared no competing interest.

