## supplemental figures, table and extended results for "Discovery of a Novel 3site State as the Multi-Substrate Bound State of P450cam"

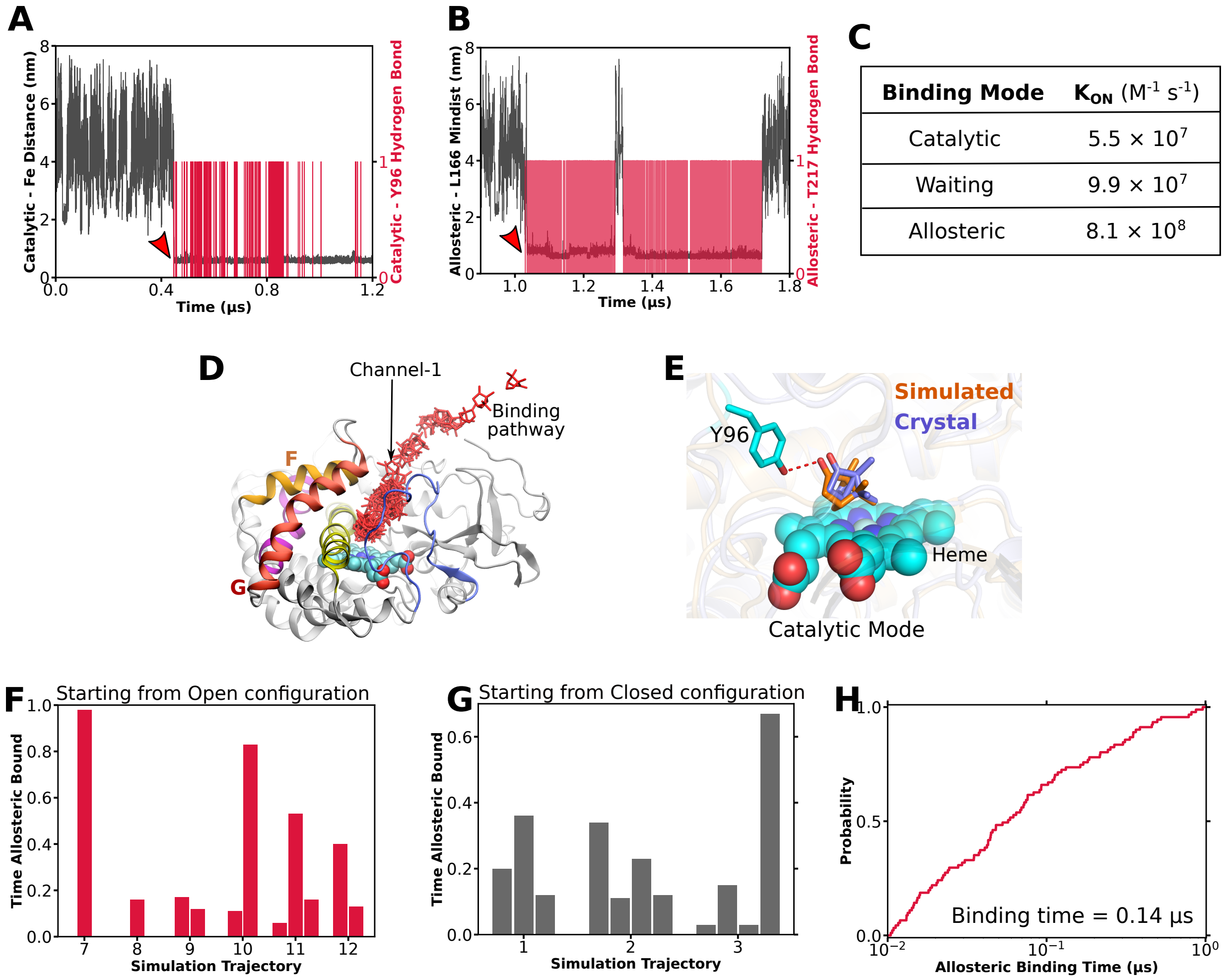


Figure S1: (A) Camphor binding in catalytic mode and hydrogen bond formation with Y96 sidechain. (B) Camphor binding in allosteric mode and hydrogen bond formation with T217. Arrowheads in (A) and (B) indicate the time of binding. (C) $k_{on}$ values of camphor binding for different binding modes. (D) Binding pathway for active site binding modes (catalytic and waiting) through channel-1. (E) Matching of catalytic binding mode (simulation) with previously known binding mode (crystal, 4JX1). (F) Fraction of total time a camphor remain bound to allosteric pocket shown for remaining binding simulations (Traj 7-12). Each bar represents a camphor molecule, such that multiple bar mean that more than one camphor molecules bound to the allosteric site (at different times). (G) Allosteric site bindings as measured in previously reported binding simulations performed with closed conformation (channel-1) of P450cam. (H) Empirical probability of binding in the allosteric site as a function of time. Mean binding time is mentioned.

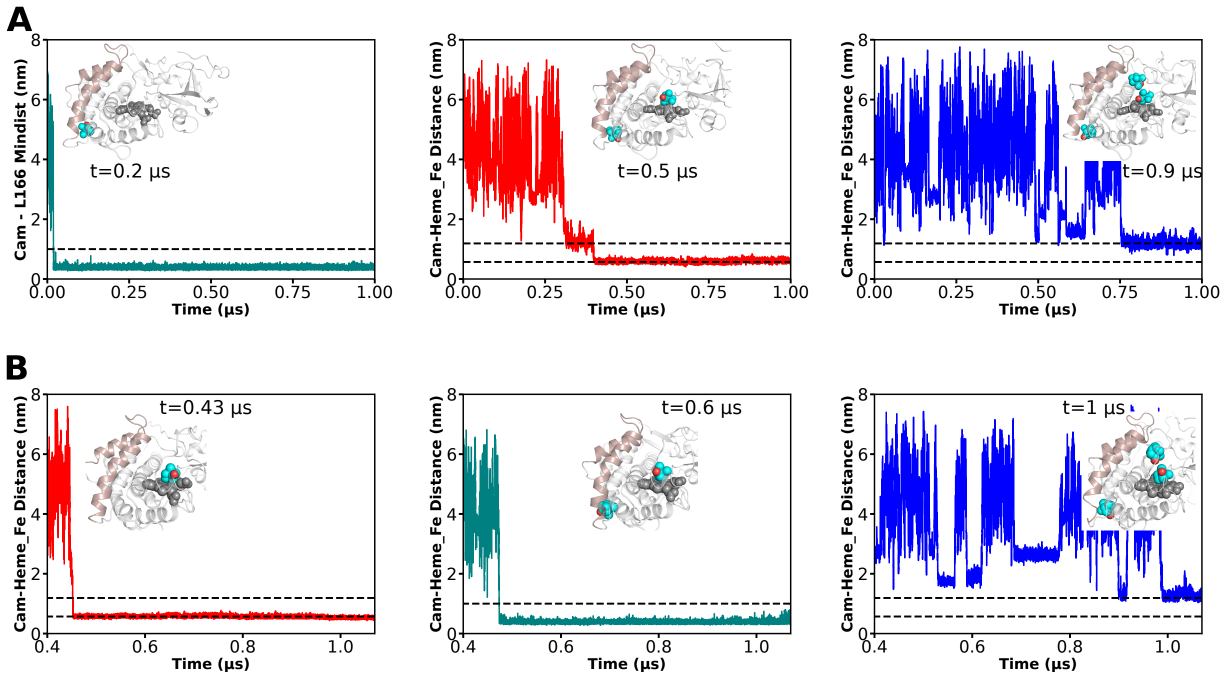


Figure S2: (A,B) Three simultaneous camphor bindings to P450cam (3site state) reproduced in other binding simulations. Color of binding curve represented the binding mode; teal-allosteric, red-catalytic, blue-waiting. Dashed line in Cam-L166 mindist represents bound cutoff for allosteric mode at 1 nm. Dashed lines in Cam-Heme_Fe distance represents the distinct locations of catalytic and waiting modes at 0.57 and 1.19 nm respectively (see Figure 2).

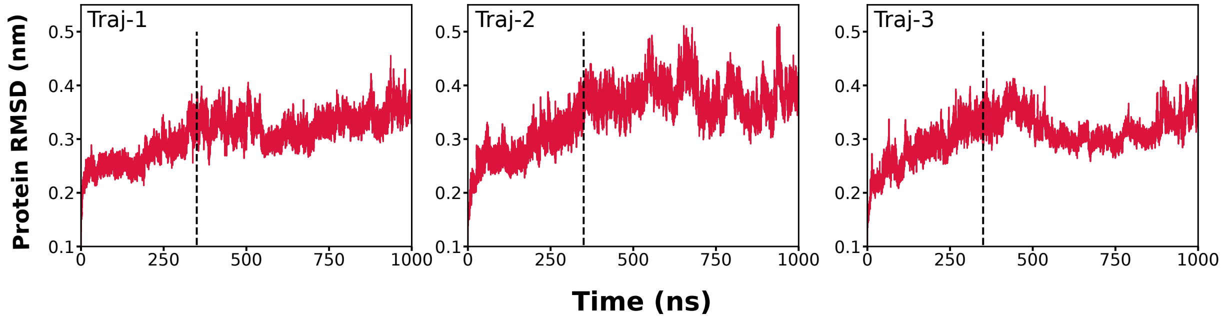


Figure S3: RMSD time profile of protein showing that system gets equilibrated within approximately 350 ns. The simulation trajectories after 350 ns were considered for analysis.

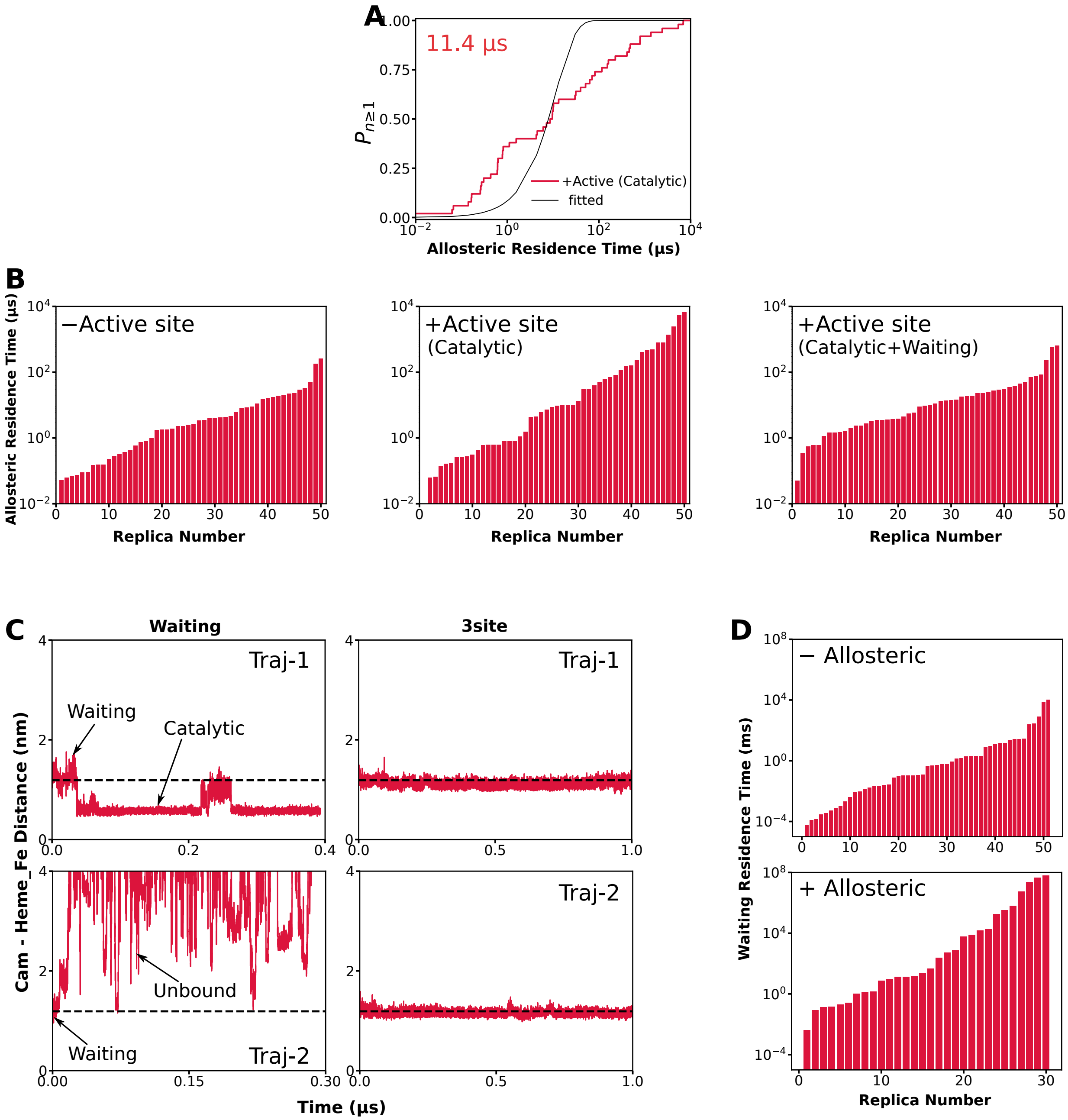


Figure S4: (A) Probability of allosteric mode unbinding with only catalytic mode in active site. Value represent the residence time. Goodness of fit is 0.012 (B) Raw allosteric unbinding times measured over different metadynamic replicas, with different state of active site. (C) Waiting binding mode stability in waiting state (unstable) and 3site state (completely stable) calculated for two different trajectories. Dashed line indicate 1.19 nm Cam-HemeFe distance representing site of waiting binding mode. (F) Raw unbinding times for waiting mode with and without the allosteric mode.

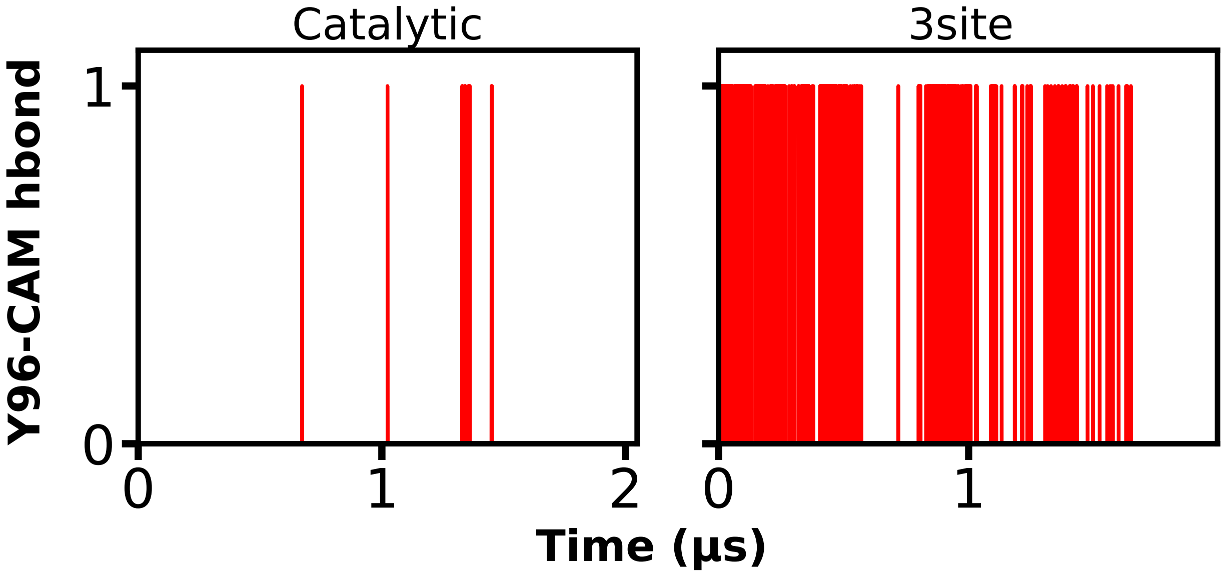


Figure S5: Hydrogen bond between substrate camphor and Y96 sidechain in different bound states of P450cam. Stable hydrogen bond formed only in 3site state.

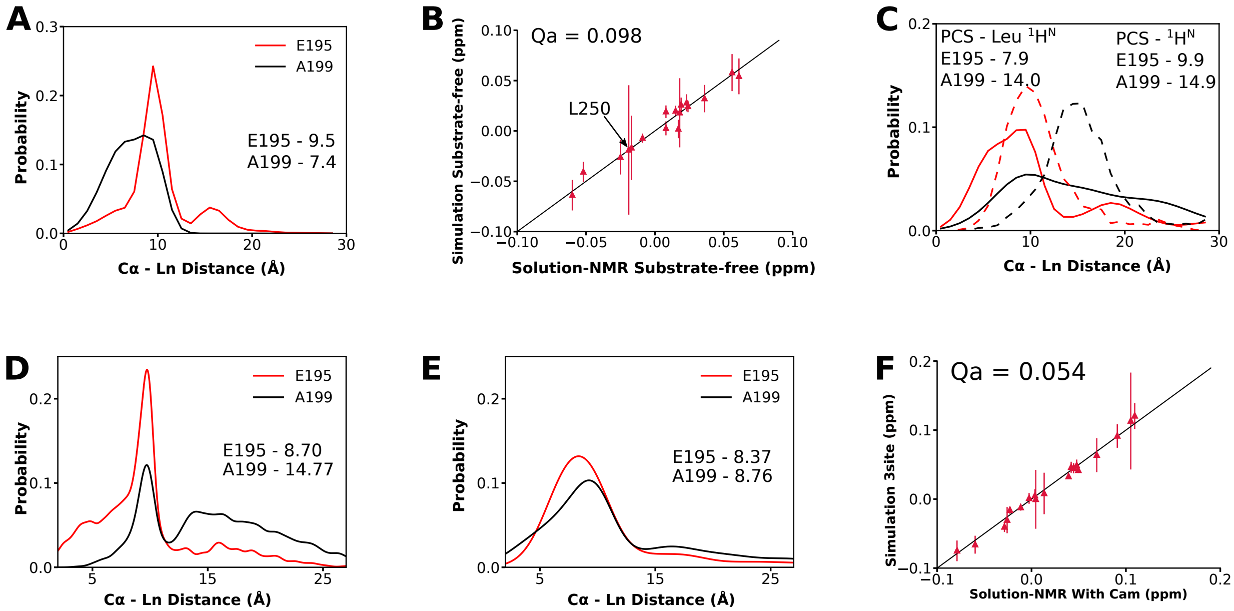


Figure S6: (A) The probability distribution of C$\alpha$-Ln distance for residues E195 (red) and A199 (black) obtained by fitting $^1H^N$ Leu PCS data (substrate-free P450cam) with substrate-free simulation ensemble. (B) PCS correlation for solution NMR and simulation PCS for substrate-free state. Large uncertainity in L250 was expected.[1] (C) The probability distribution of C$\alpha$-Ln distance for residues E195 (red) and A199 (black) obtained by fitting PCS data (P450cam+Cam) with substrate-free simulation ensemble. Solid lines represents $^1H^N$ Leu PCS while dashed line represents $^1H^N$ PCS both measured in presence of substrate camphor. (D,E) The probability distribution of C$\alpha$-Ln distance for residues E195 (red) and A199 (black) obtained by fitting $^1H^N$ Leu PCS data (P450cam+Cam) with catalytic (D) and 3site (E) simulation ensmeble. (F) PCS correlation of solution NMR $^1H^N$ Leu PCS and simulation PCS for 3site state. **Note**: Straight line in B,F represents perfect correlation. Numerical values in A,C,D,E represents the fitted mean values of distance distributions.

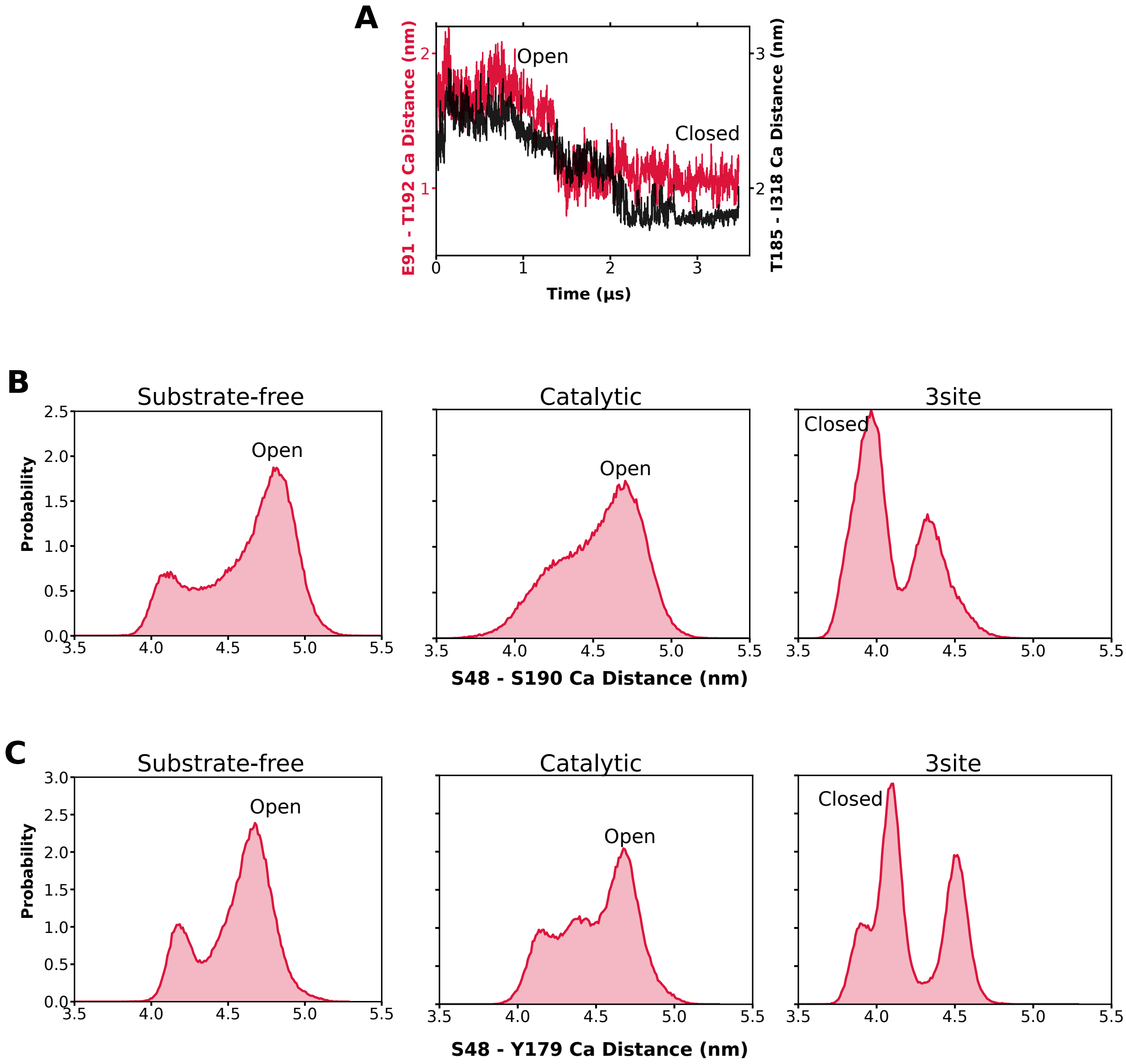


Figure S7: (A) Channel-1 conformation in binding simulation involving 3 simultaneous bindings i.e., 3site model. Starting from open conformation, channel-1 transitions to closed conformation in real time. (B,C) Channel-1 open/closed conformations for different states of P450cam as measured by probability distribution of S48-S190 or S48-Y179 Ca distance. Dominant closed conformation of channel-1 was observed only in 3site state.

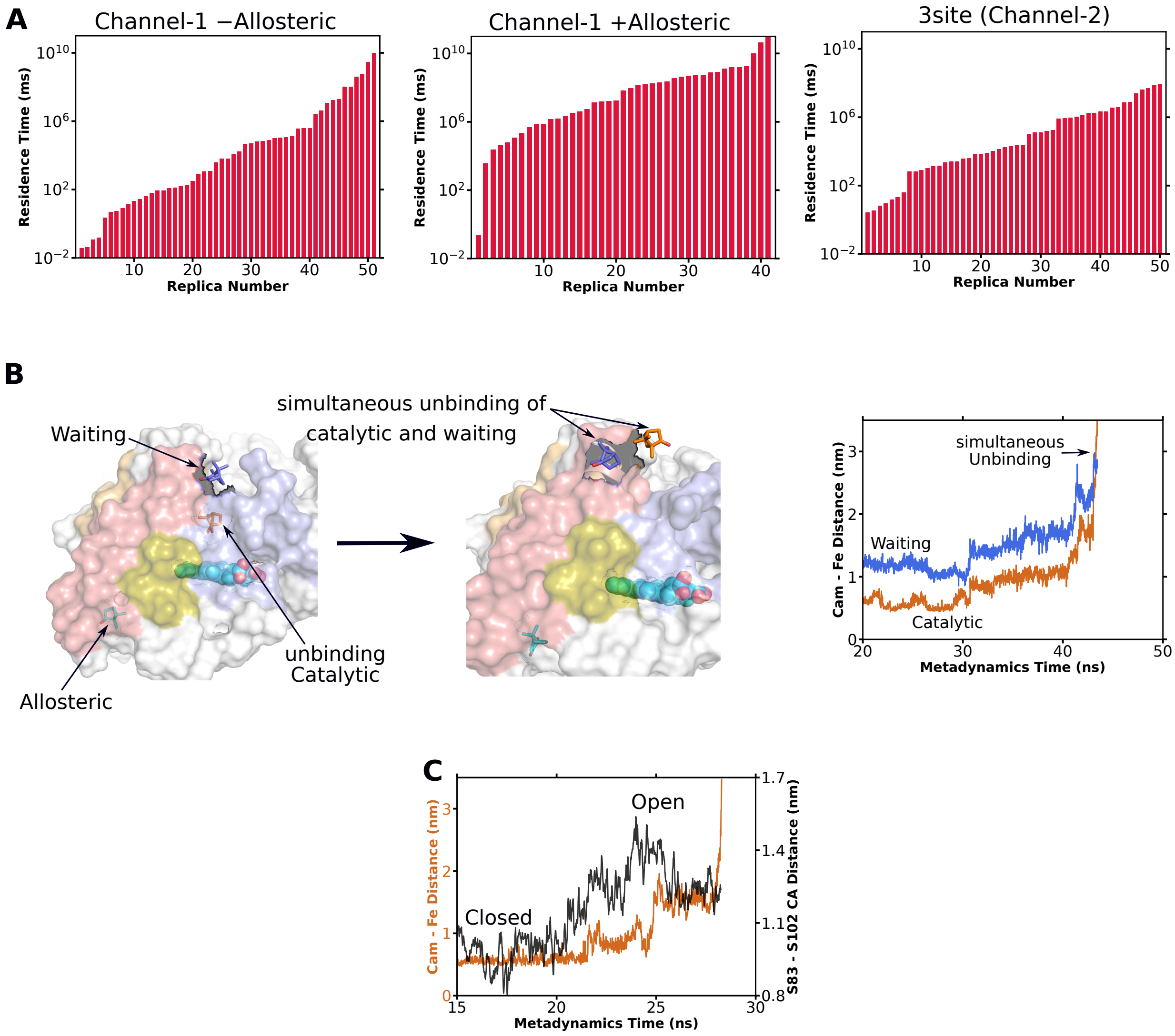


Figure S8: (A) Raw unbinding times for catalytic mode through different channels. (B) Simulataneous unbinding of catalytic and waiting modes through channel-1. Unbinding catalytic through channel-1 pushed on the waiting mode to unbind. (C) Opening of channel-2 with catalytic mode unbinding.

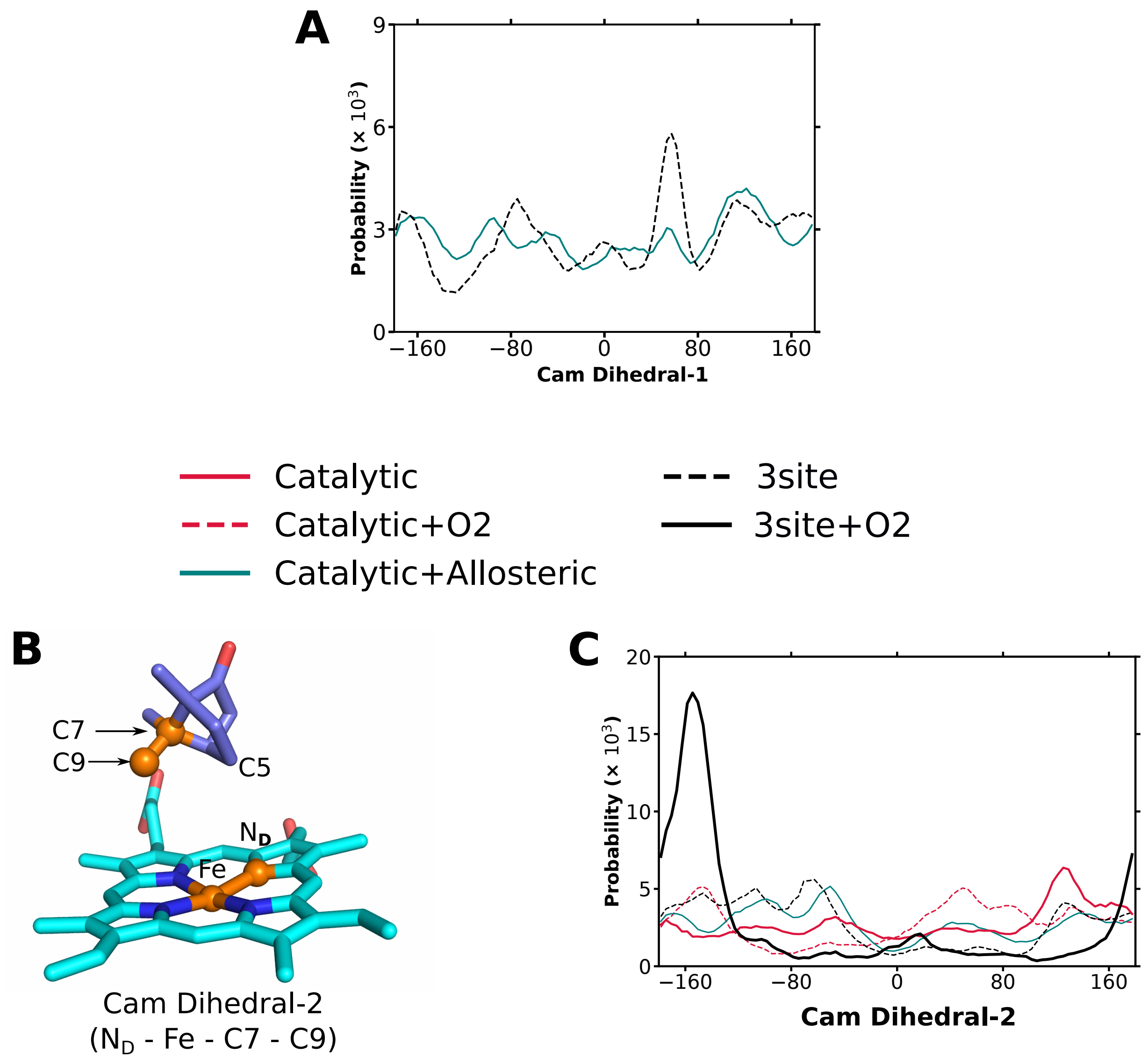


Figure S9: (C) Probability distribution of cam Dihedral-1 in different states of P450cam. (B) Catalytic binding mode depicting cam Dihedral-2. (C) Probability distribution of cam Dihedral-2 in different states of P450cam.

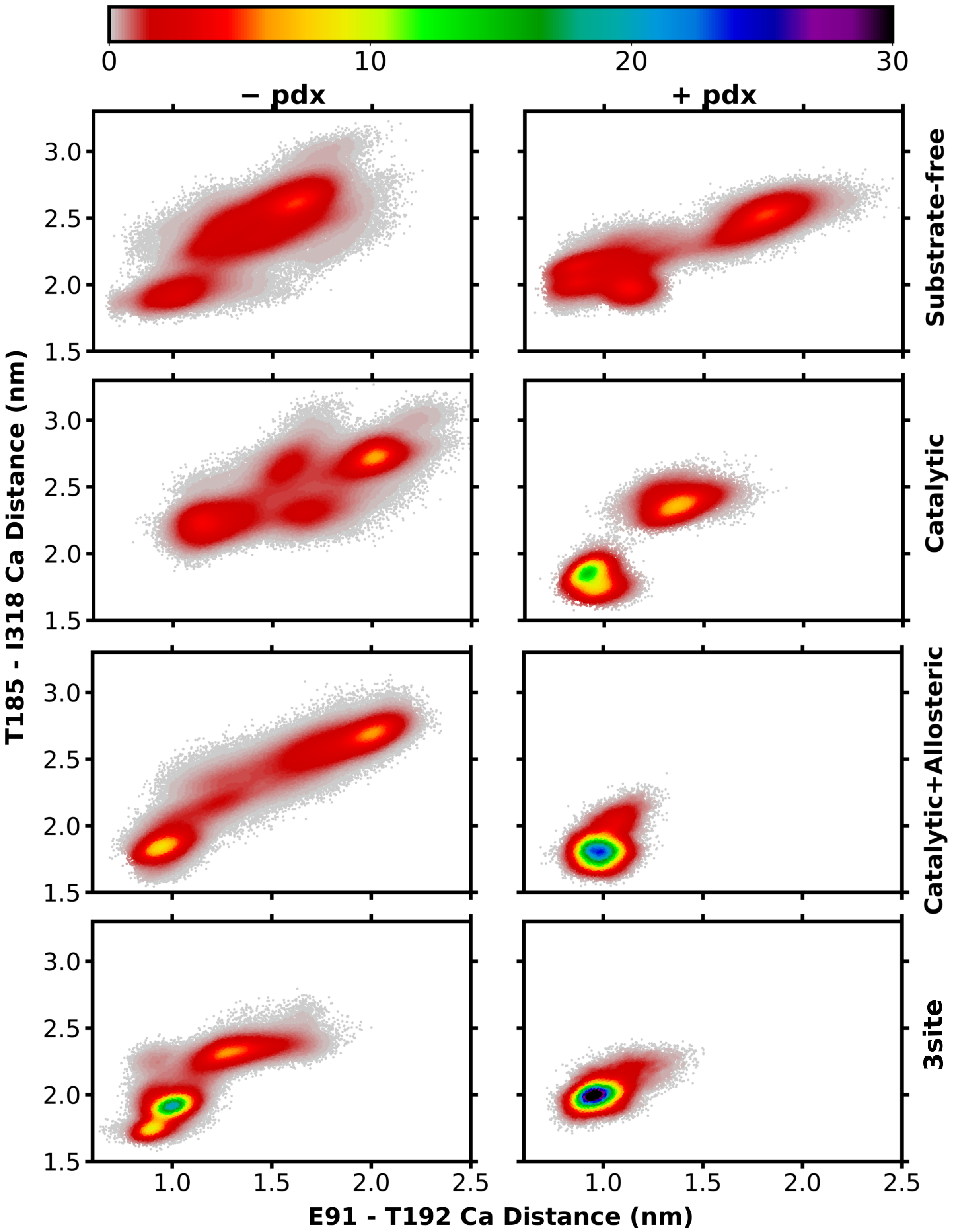


Figure S10: One to one comparison of channel-1 (probability density distribution) with and without Pdx over different states of P450cam.

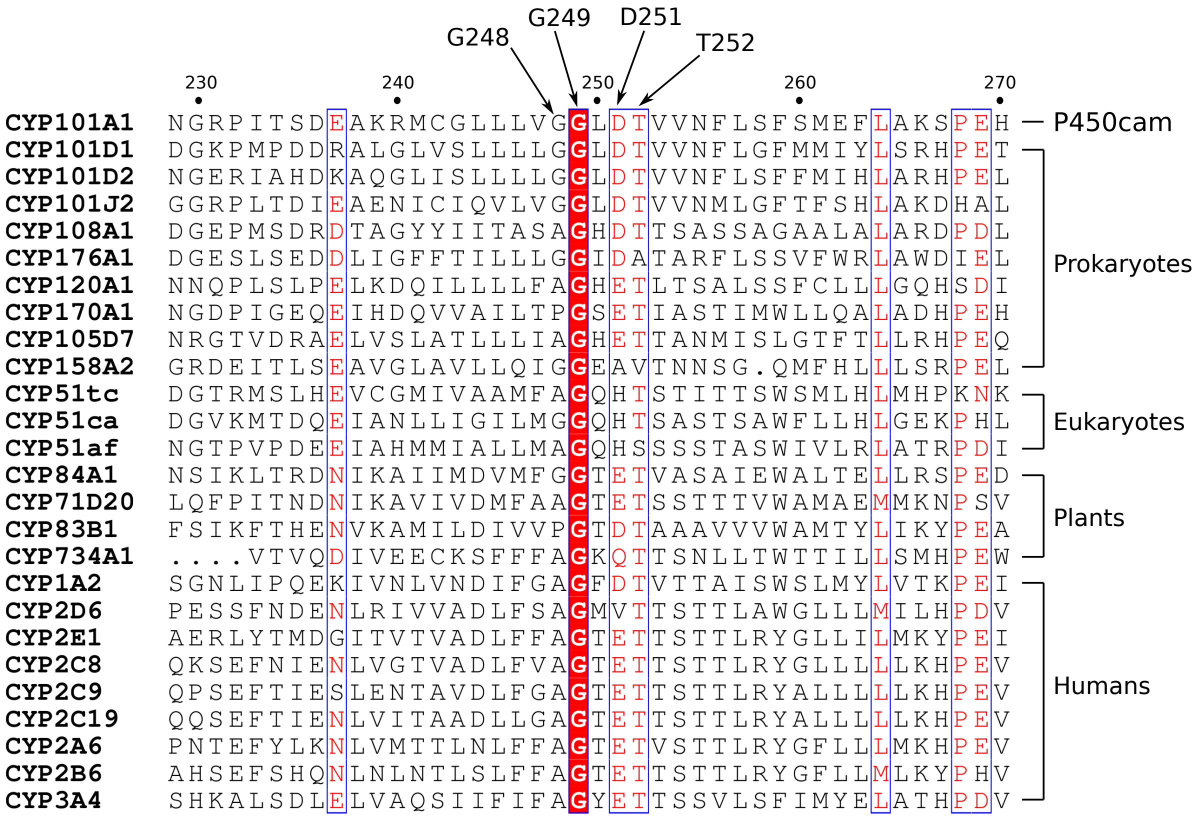


Figure S11: Evolutionary conservation of kink generating glycine residues (G248, G249) of I helix as detected by multiple sequence alignment over different species ranging from prokaryotes to humans.

### Supplementary Methods

**SM-1: Simulation ensembles of different states of P450cam**: Separate simulation ensembles were generated for different states of P450cam binding process namely substrate-free, catalytic, allosteric, waiting, catalytic+allosteric and 3site (catalytic+allosteric+waiting) states (Figure 3, main text). For each state, P450cam in open conformation (chain A of 4JX1) was used. The camphor molecule(s) were circumspectly placed in the desired binding mode(s) based on the knowledge from unbiased binding simulations. For instance, for catalytic, allosteric and waiting states, one camphor molecule was placed in the catalytic, allosteric and waiting binding modes by maintaining the camphor location and orientation based on the knowledge from binding simulations (results). Similarly, for catalytic+allosteric states, two camphor molecules were placed in catalytic and allosteric binding modes while for 3site state, three camphor molecules were placed at catalytic, waiting and allosteric binding modes. The P450cam along with bound camphor molecule(s) (only P450cam in case of substrate-free) was placed at center of cubic box and filled with TIP3 water molecules and $Na^+$ ions (to neutralize) the system. The protonation states of amino acids, force field parametrization and system equilibration was done same as for binding simulations. For each states, at least 3 replicates of long unbiased simulations were performed, each of 1 $\mu$s time length. The simulations were stopped prematurely, if the camphor molecule was found to be unstable in its binding mode, for instance in waiting state (results).

For simulation ensembles of Pdx bound P450cam states (Figure S13A-E), starting structures of substrate-free, catalytic, allosteric, catalytic+allosteric and 3site states were used. In addition to P450cam and camphor, the starting structures also contained Pdx (residue 1-106) bound to P450cam proximal surface based on crystallographic pose of 4JX1[2] (corresponding to chain A and C). The open P450cam + Pdx + camphor(s) (not in substrate-free) structure was placed at center of cubic box and filled with water and $Na^+$ ions. The forcefield parametrisation and system equilibration was same as for binding simulations. Additionally, Pdx contained an iron sulfur cluster ($Fe_2S_2$) coordinated with 4 cysteine residues. The forcefield parameters for coordinated cysteines and $Fe_2S_2$ were generated using ab-initio QM simulation.(See below) For each state, at least 3 replicates of unbiased MD simulations were performed, each corresponding to a maximum of 1 $\mu$s. In some simulations, the Pdx was found to be unstable, where only the stable portions of the simulations were considered for further analysis. Overall, simulation ensembles of Pdx were smaller as compared to non-Pdx counterparts. Nevertheless, these are the longest simulations of P450cam+Pdx complex reported till date.

For simulation ensembles of catalytic+$O_2$ and 3site+$O_2$ states (Figure S13F,G), systems were generated exactly as for catalytic and 3site states. But instead of heme, oxy-heme ($O_2$ molecule coordinated Fe atom of heme) was used. The oxy-heme was parametrised according to CHARMM27 forcefield.[3] Heme parameters of CHARMM27 were exactly same as for heme in charmm36, only two oxygen atoms were additionally present coordinated to Fe atom of Heme. For mutant states, i.e., L166A-substrate-free, L166A-catalytic, L166A-allosteric, L166A-catalytic+allosteric and T217V-substrate-free (Figure S13H-L), systems were generated exactly as their wildtype counterparts, but the P450cam was mutated at 166 from leucine to alanine or at 217 from threonine to valine. The mutations were performed in-silico using charmm-gui.[4] Forcefield parametrization and system generation was same as for other states of P450cam and unbiased MD simulations were performed for a maximum time of 1 $\mu$s for 3 replicates, unless mentioned otherwise. For L166A-allosteric and L166A-catalytic+allosteric states, the allosteric mode was highly unstable (results), hence the simulations were stopped prematurely as soon as the allosteric ligand unbind spontaneously. Additionally, simulations of T217V-allosteric were performed only for 400 ns and were only used for extracting frames for metadynamic simulations and were not used for any analysis.

**SM-2: Forcefield parameterization of iron-sulfur cluster in Pdx**: The redox partner of P450cam i.e., putidaredoxin (pdx) contained an iron-sulfur cluster of type $[Fe_2S_2(CYS)_4]^{2-}$, for which the CHARMM36 compatible forcefield parameters were generated using ab-initio quantum mechanics (QM) simulations. The starting configuration of iron-sulfur cluster along with 4 coordinated cysteines (C39, C45, C48 and C86) was taken from published pdx structure (corresponding to chain C of PDB id - 4JX1; Figure S14). To satisfy the valencies, 'H' and 'OH' moities were added to amide and carbonyl terminals of coordinated cysteine residues. Calculated spin densities with guess wave function indicated the oppositely coupled spins of two Fe centers, hence leading to net zero spin and multiplicity of one. Geometry optimization of the starting configuration was performed using Density Functional Theory (DFT[5]) using Gaussian-16[6] suite of programs. Calculations were performed using local meta-General Gradient Approximate (GGA) exchange correlation functional M06L[7] with 6-31G**[8] and SDD(ECP)[9] as basis set for main group and transition metal atoms respectively. Charges from electrostatic potentials using grid based method was employed to calculate the ESP[10] charges. While performing the CHELPG calculations, radii of 2.05 $\AA$ was specified for Fe atom. Harmonic frequencies were calculated for optimized structure and were identified with no imaginary frequencies.

The gaussian-16 generated log and fchk files were used to generate the forcefield parameters for iron-sulfur cluster. The potential energy function of CHARMM36 forcefield is constituted of bonded and non-bonded terms, i.e., bonds, angles and dihedrals as bonded terms, while partial atomic charges and Lennard-Jones parameters as non-bonded terms. The equilibrium values and force constants for bond and angle terms were generated by QM acquired geometric and vibrational harmonic frequencies using modified seminaro approach[11] with scaling factor of 2.8. For dihedrals, which could not be generated by seminaro approach, the force constants were attributed with the value of 0.8 kJ/mol, the Tohiko Ichiye[12] value for iron-sulfur clusters. The equilibrium values for dihedrals were attributed with $0^o$ to maintain the planarity of $Fe_2S_2$ molecule. The multiplicity of dihedrals was provided as per the possible rotameric states. The existent parameters of CHARMM36 forcefield were used for Lennard-Jones terms. The partial atomic charges were attributed with ESP charges. Finally, the CHARMM36 forcefield folder was updated to included parameters for $[Fe_2S_2(CYS)_4]^{2-}$ as composed of two molecules, i.e., CSG (cysteine molecule without gamma sulfur (Sg), similar to CYS3 molecule of Dr. Da Silva[13]) and FES molecule ($Fe_2S_2Sg_4$). The updated forcefield is provided in supplementary, along with README files to track the changes/additions. All the changes in the forcefield can be tracked by searching for keyword JM.

Ultimately, the reliability of updated CHARMM36 forcefield was confirmed by performing long unbiased MD simulations of Pdx protein. Pdx along with iron-sulfur cluster was taken in center of cubic box and filled with water molecules and $Na^+$ ions. Fe atoms of iron-sulfur cluster were coordinated to gamma sulfur atoms of four cysteine residues, namely C39, C45, C48 and C86. The four coordinated cyteines were modelled as above mentioned CSG while rest as normal cysteines. Overall, the system contained 19795 atoms and parameterized by updated CHARMM36 forcefield. Firstly, followed by 5 ns NVT and NPT equilibrations, a 100 ns simulation was performed. ESP Charges and energies were recalculated by QM simulations for 100 iterations with structures taken after every 1 ns from aforementioned 100 ns simulations. The mode ESP charges from 100 iterations were used in final updated forcefield. Ultimately, 3 independent production runs were performed each of 1 us time length with the final updated forcefield. Last 650 ns of each simulation were combined as final simulation ensemble of Pdx, which was analyzed further. All the bond lengths, angles and dihedrals were measured from the Pdx simulation ensemble, it was found that all the parameters were stable and followed the unimodal gaussian distribution (indicative of stable configuration, Figure S14). The fitted gaussian mean values perfectly resembles the updated forcefield values and reported crystal configuration of I. F. Sevrioukova.[14] The dihedrals deviated slightly from planarity with +- 11 degrees (Figure S14). The fitted standard values were used as measure of error/spread and were found to be significantly low. Additionally, forcefield parameters of iron-sulfur cluster in partially reduced form i.e., $[Fe_2S_2(CYS)_4]^{3-}$ were also generated with the above procedure and all parameters were found stable and closely similar to above state.

**SM-3: Free energy calculations**: For calculating free energy change for binding of catalytic binding mode, the umbrella sampling approach was used. The binding at catalytic mode was measured using center of mass distance between camphor and Heme-Fe and hence used as a collective variable (CV) in umbrella sampling.[15] The value of CV ranged from 0.45 to 2.0 nm representing bound and unbound extremes of catalytic mode. The CV range was discretized into 32 windows with an uniform interval of 0.05 nm between consecutive windows. The free energy of catalytic mode was mapped in two states i.e., with and without allosteric binding mode. For both the states, the starting structures corresponding to each umbrella were taken from binding simulations. For 'without allosteric mode', the starting structures were choosen such that allosteric site was not occupied by any camphor molecules, while for 'with allosteric mode', starting structures were choosen such that allosteric site was stably bound with substrate camphor. Each umbrella window was equilibrated for NVT and NPT and finally sampled for 25 ns ensuring sufficient sampling of entire CV range by restraining each umbrella to desired value of CV using suitable harmonic restraint potential. Finally, weighted histogram analysis method (WHAM)[16] was used to reweight the umbrellas to get free energy of catalytic binding mode as a function of distance between catalytic camphor and Heme-Fe. The error was estimated using bootstrapping over 1000 iterations. The binding free energy was calculated as difference between free energy of bound (the minima of curve) and unbound (at 2.0 nm) states and corrected for standard state.

The umbrella sampling approach was also used for mapping the underlying thermodynamics of channel-2 opening-closing transition. The opening of channel-2 was measured by distance between C$\alpha$ atoms of S83 and S102 and used as CV for umbrella sampling calculations. The S83-S102 distance ranged between 0.45 to 1.65 nm representing the open and closed extremes of channel-2. The CV was descretized into 25 windows with an uniform interval of 0.05 nm between consecutive umbrellas. The channel-2 thermodynamics were measured for 5 different states i.e., in absence of camphor (substrate-free), in presence of catalytic, or allosteric or catalytic+allosteric or catalytic+waiting+allosteric (3site) camphor molecules. The starting structures were taken from simulation ensembles of corresponding state. The simulations were performed similarly as described in above section and underlying free energy curve was calculated as a function of CV.

For free energy of binding of allosteric mode, the probability density distribution of camphor location in binding simulations was constructed as a function of minimum distance between heavy atoms of camphor and L166 (CV). All the 4 camphor molecules of all 12 binding simulations were considered for calculation to ensure extensive sampling of CV. The standard binding free energy was calculated as per eq(1):

$$\Delta G^o = k_B T \cdot ln[\frac{P(B)}{P(U)} \times \frac{V_o}{V}] \qquad (1)$$

where: $k_B$T = 0.6 kcal/mol, P(B) and P(U) are probability distributions of bound (less than 1 nm) and unbound states, $V_o$ = 1661 $\AA^3$ and V is accessible volume for camphor in binding simulaitons.

**SM-4: Residence time calculation by unbinding simulations**: Well tempered infrequent metadynamics[17] approach was utilized to calculate the residence time of camphor in different binding modes. This novel method allows accurate measurement of residence time of ligand within any binding pocket.[18] In this method, bias was periodically applied (slower than bottleneck times) to bound ligand as a function of CV to propel the ligand to unbind.[19] Note that, in case of this study including multiple binding modes, the bias was applied only on one particular binding mode (for which the residence time was being calculated) while not on others (if present). The detailed description is as below.

For catalytic binding mode, the residence time was calculated in 3 different states i.e., in absence of other binding modes (catalytic state), in presence of allosteric mode (catalytic+allosteric state), in presence of allosteric and waiting modes (3site state). In catalytic and catalytic+allosteric states, the catalytic mode was biased to unbound only through channel-1 (results). This was achieved by using distance between Heme-Fe and camphor as CV. For each state, the starting structures were taken from the simulation ensembles of corresponding state. The 50 structures corresponding the lowest values of CV in the simulation ensemble were taken, hence representing the stable most bound poses. Starting from a stable bound pose, a history dependent bias potential [V(s,t)] was periodically applied to catalytic mode as a function of CV. The bias was applied in the form of gaussians of height 1.2 kJ/mol and width 0.035 nm and periodically added after every 5000 steps (corresponding to 10 ps). The height of gaussians were tempered by biasing factor of 6. The metadynamics simulations were continued until the catalytic mode reached the unbound state (camphor heme-Fe distance greater than 3 nm, representing the camphor distinctly away from P450cam). The bias potential added to unbind the catalytic mode was used to calculate the acceleration factor ($\alpha$) as per eq(2). The unbiased unbinding time ($T_U$, or transition times) was calculated as product of acceleration factor and metadynamics time ($T_M$, time of unbinding observed in metadynamics simulation) as per eq(3). 50 such unbiased transition times were calculated corresponding to each starting structure. The obtained transition time statistics was subjected to Poisson analysis because the unbiased ligand transitions times from bound to unbound state follows the poisson distribution.[20] The empirical cumulative distribution function (ECDF) calculated from 50 unbiased transition times was fitted with cumulative distribution function (CDF) of poisson distribution given in eq(4) and the residence time ($\tau$) was estimated by curve fitting. For error analysis, the goodness of fit was confirmed using 2 sample Kolmogorov-Smirnov (KS) test[21] and fitting greater than 0.01 was considered good.

$$\alpha = \langle e^{\beta V(s,t)} \rangle_{T_M} \tag{2}$$

$$T_U = T_M \times \alpha \tag{3}$$

$$P_{n\geq 1} = 1 - exp(-\frac{T_M}{\tau}) \qquad (4)$$

where;

V(s,t) - bias potential as a function of CV (s) and time (t)

$\beta = \frac{1}{k_B T}$ - inverse temperature (T), $k_B$ is Boltzman constant

$P_{n\geq 1}$ - probability of observing atleast one transition within time $T_M$

For residence time calculation of catalytic mode in 3site state, the 50 starting structures were taken from 3site simulation ensemble corresponding to stable bound pose. The bias potential was applied on catalytic mode to propel it out from catalytic center. The distance between camphor and catalytic center (i.e., Heme-Fe, catalytic residues D251, T252 and pocket kink residue V396) was used as CV such that catalytic mode was free to unbind through any possible path (channel-1 or channel-2, results), although presence of waiting mode restricts unbinding through channel-1 to some extent. The bias potential addition and residence time calculation was performed exactly as described above for catalytic mode.

For waiting binding mode, the residence time was calculated for 2 different states, i.e., with and without allosteric binding mode. Since, waiting mode is stable only in 3site state, the metadynamics simulations performed for unbinding catalytic mode through channel-1 were used for estimating the residence time of waiting mode. While the catalytic mode unbound through channel-1 (based on distance between camphor and Heme-Fe), it goes through waiting state. Hence, transition times for waiting mode were measured as catalytic mode transitions from waiting state (CV = 1.2 nm) to unbound state (CV $\geq$ 3 nm). Further residence time calculations were performed as described above in eq(2-4).

For allosteric binding mode, the residence time was calculated in 4 different states, i.e., in absence of other binding modes (allosteric state), in absence of other binding modes but with T217V mutation (T217V-allosteric state), in presence of catalytic mode (catalytic+allosteric state) and in presence of catalytic and waiting modes (3site state). The 50 starting structures were taken from simulation ensembles of the corresponding states representing the stable allosteric bound poses. The bias potential was added to allosteric mode only, as a function of minimum distance between heavy atoms of camphor and L166. The gaussian bias of height 0.4 kJ/mol and width 0.035 nm was added periodically after every 5000 steps (10 ps) with a tempering factor of 4. The metadynamics simulations were continued until the allosteric mode leaved the allosteric site. The unbinding was defined with minimum distance between camphor and L166 of greater than equal 1 nm as observed in free energy of allosteric mode binding. The unbiased transition times and residence times were calculated as described above using eq(2-4) and goodness of fit was checked by 2 sample KS test with statistic greater than 0.05.

Additionally, residence time of allosteric mode was calculated for L166A mutant. In this case, the allosteric mode was found to be very unstable such that camphor was spontaneously leaving allosteric site within hundred of ns of unbiased simulation. Hence for L166A mutant, the unbiased binding times were taken from unbiased simulations only. For instance, the unbiased simulations were started with camphor in allosteric mode (both with and without active site modes) and the simulation was continued till the allosteric mode spontaneously leave the allosteric site. Hence, no bias was added. A total of 58 such simulations were performed. The residence time was calculated as per eq(4) by curve fitting and KS test as describe above.

Note that the fit between ECDF of transition times and CDF of poisson was almost perfect (very high KS statistic) in most cases except for some cases of catalytic mode. Additionally, not all trajectory result in unbinding in catalytic+allosteric state while calculating residence time of catalytic mode. In this case, to FIT the ECDF of transition times and CDF of poisson, some outliers were removed and KS statistic of 0.01 was used. Nevertheless for every residence time calculation, along with the poisson statistics the raw data for all transition times were provided in supplimentary.

**SM-5: Pocket Analysis**: The pocket analysis was performed to calculate the 3D volume and constituting residues for active and allosteric sites. The pocket volume of active site was measured on starting substrate-free configuration of binding simulations (corresponding to chain A of 4JX1). For allosteric site, the pocket analysis was performed on aligned structures taken after every 1 ns from the simulation ensembles i.e., substrate-free, catalytic, allosteric, catalytic+allosteric, 3site, L166A-substrate-free and L166A-catalytic states. The MDpocket[22] utility was used to sample the pocket architecture using Voronoi tessellation algorithm.[23] The pocket volume was calculated as cumulative volume of alpha spheres detected in a particular site (allosteric site) for more than 50% of simulation ensemble. In some frames ( 4-5%) the pocket cannot be detected either due to sidechain flexibility or limitation of MDpocket, such frames were not considered. The pocket volumes of detected frames was bootstrapped for 10000 iterations for smoothing of probability density curve (mean remain unchanged). Further for each frame, list of atoms in contact with alpha spheres of pocket were derived from MDpocket logfile. The fraction of $i^{th}$ amino acid ($M_i$) constituting the allosteric pocket was calculated as weighted sum of its atoms in contact with alpha spheres and normalized by total weighted sum and number of frames as per eq(5). The atoms were weighted into three categories i.e., polar (0.5), apolar (0.35) and hydrogens (0.15).

$$M_i = \langle \frac{\sum a_j w_j}{\sum a_k w_k} \rangle_n \qquad (5)$$

where a - atom type; w - weight; n - number of simulation frames; j - sum over atoms in contact with alpha spheres; k - sum over all atoms of an amino acid

**SM-6: Channel-1 order parameters**: Channel-1, an opening near the junction of F, G and $\beta'$ helices which upon opening allowed direct access to active site, is the most studied aspect of P450cam. The opening of channel-1 involved retraction of F and G helices away from $\beta'$ and/or simultaneous unfolding of $\beta'$ helix.[2,24] It has been postulated that instead of simple retraction of F and G helices, channel-1 opening is a multifaceted conformational change and F and G helices can behave independently.[25] Additionally, $\beta'$ helix can undergo unfolding and displaced away from F/G loop. Although, several CVs have already been defined, here we devised two distance based CVs that can suitably detect the channel-1 opening by considering most of its aspects. First, distance between C$\alpha$ atoms of T192 (G helix) and E91 ($\beta'$ helix) to detect the separation of G and $\beta'$ helices away from each other and unfolding of $\beta'$; second the distance between C$\alpha$ atoms of T185 (F helix) and I318 (318$\beta$, a stable reference secondary structure) to detect the retraction of F helix (Figure S15A). Overall, these two distance based CVs have reliably detected the simultaneous yet independent retraction motions of F and G helices (Figure S15B) as well as unfolding/displaced $\beta'$ helix. E91-T192 of 0.9 nm and T185-I318 of 1.75 nm represented the perfectly closed state as observed in simulations and crystal structures, while larger values depict the opening. Additionally, distance between C$\alpha$ atoms of S48 - S190 and S48 - Y179 were also used and represented the closest possible analogues of distances measured in previously reported DEER studies.[25,26]

**SM-7: Contribution of allosteric mode to function**: The three binding modes of P450cam contribute additively to conformational changes within the active site (results) as the P450cam goes from

$$\text{Substrate-free} \rightarrow \text{Catalytic} \rightarrow \text{Catalytic+Allosteric} \rightarrow \text{3site}$$

states. The contribution of allosteric binding mode ($P_A$) to a particular conformational change (P) within active site was defined as change in P upon allosteric binding (catalytic to catalytic+allosteric transition) normalized by total change in P (substrate-free to 3site transition) as per eq(6). The allosteric contribution was calculated for 4 different conformational changes within the active site, which were deemed important for function namely, Y96-$\beta$ dihedral, catalytic-Y96 hydrogen bond, pocket-kink and $I^{th}$ kink. Y96-$\beta$ dihedral, pocket-kink and $I^{th}$ kink exhibit bimodal distribution where two peaks in probability distribution represented active and inactive conformation. $P_i$ (i - state, eg $P_{3site}$) in eq(7) represented the fraction of active conformations for conformational change P and calculated as area under the probability distribution curve within active range. For Y96-$\beta$ dihedral, the dihedral angle of less than -120$^o$ was defined as active. Similarly D251-I395/V396 distance of less than 1 nm and $I^{th}$-318$\beta$ pairwise distance of less than 2.75 nm was used as active range for pocket kink and $I^{th}$ kink respectively. For catalytic-Y96 hydrogen bond, $P_i$ was defined as probability of hydrogen bond.

$$P_A = \frac{P_{Catalytic+Allosteric} - P_{Catalytic}}{P_{3site} - P_{Substrate-free}} \times 100 \tag{6}$$

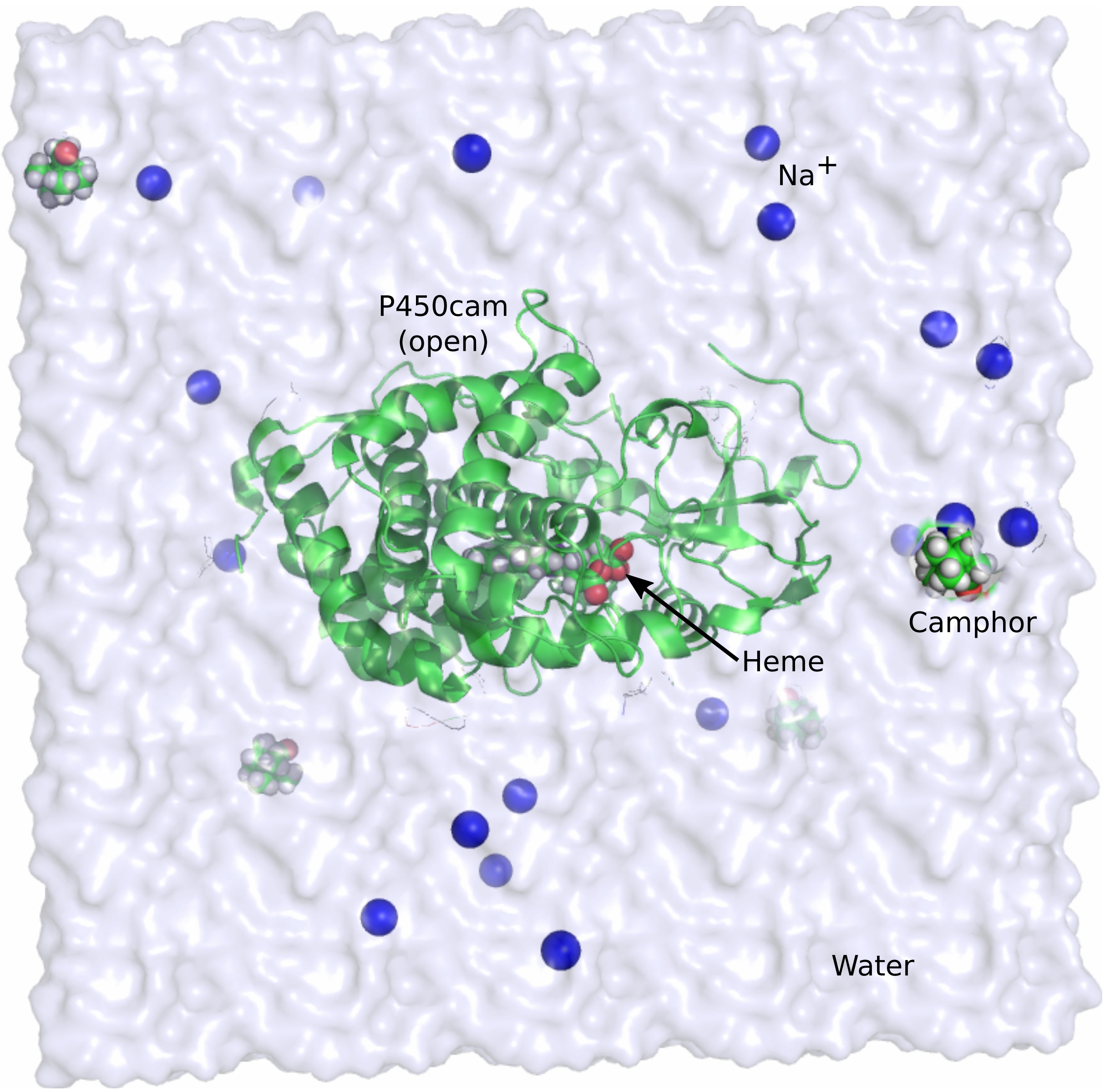


Figure S12: Starting configuration for unbiased binding simulations, consisting of open P450cam (green cartoon) at center of cubic box filled with TIP3 water (lightblue) and $Na^+$ ions (blue spheres). 4 substrate camphor molecules (green VDW) were also present in solvent, away from P450cam representing the unbound state.

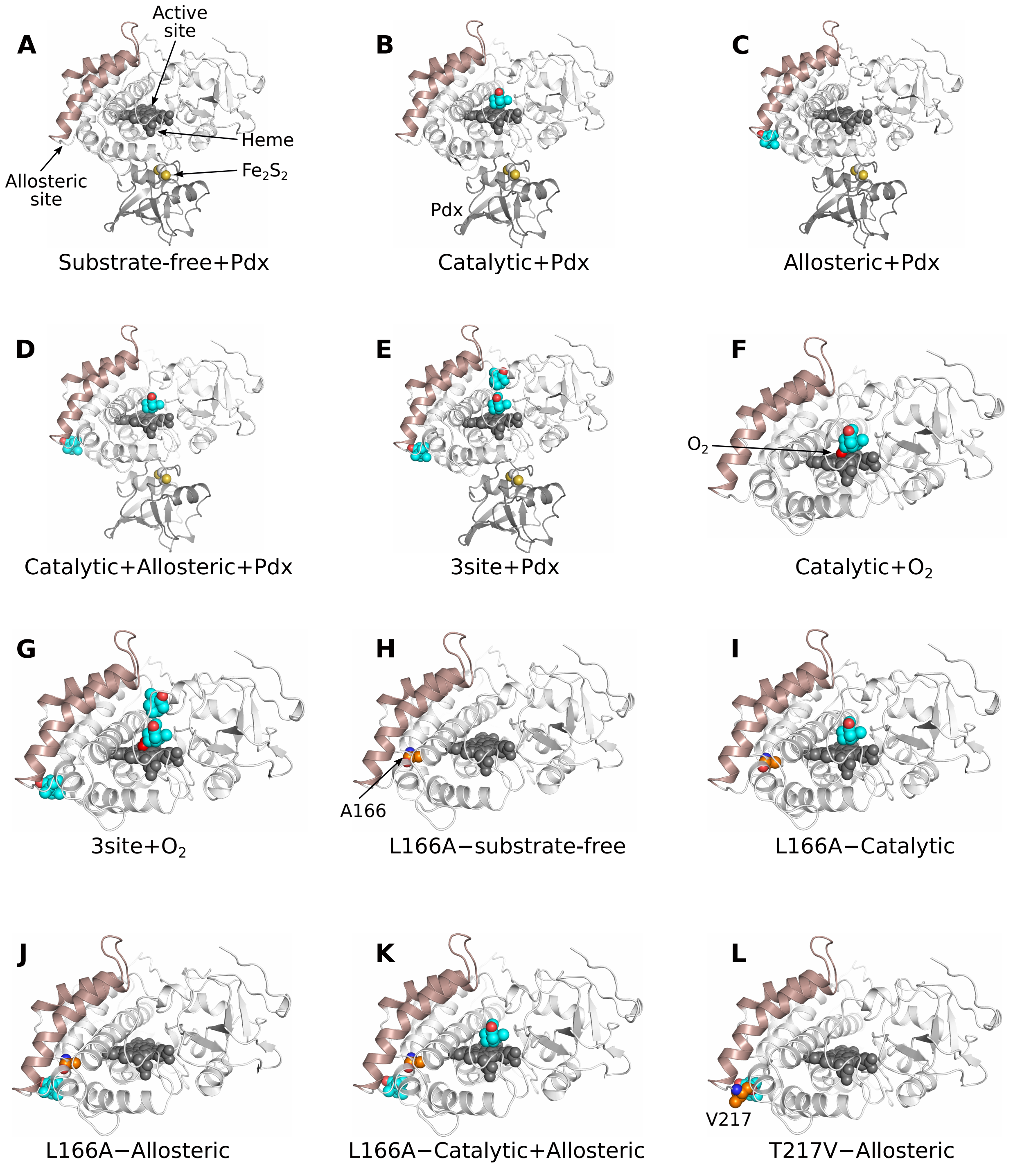


Figure S13: (A-L) Starting configurations of different states of P450cam and nomenclature used.

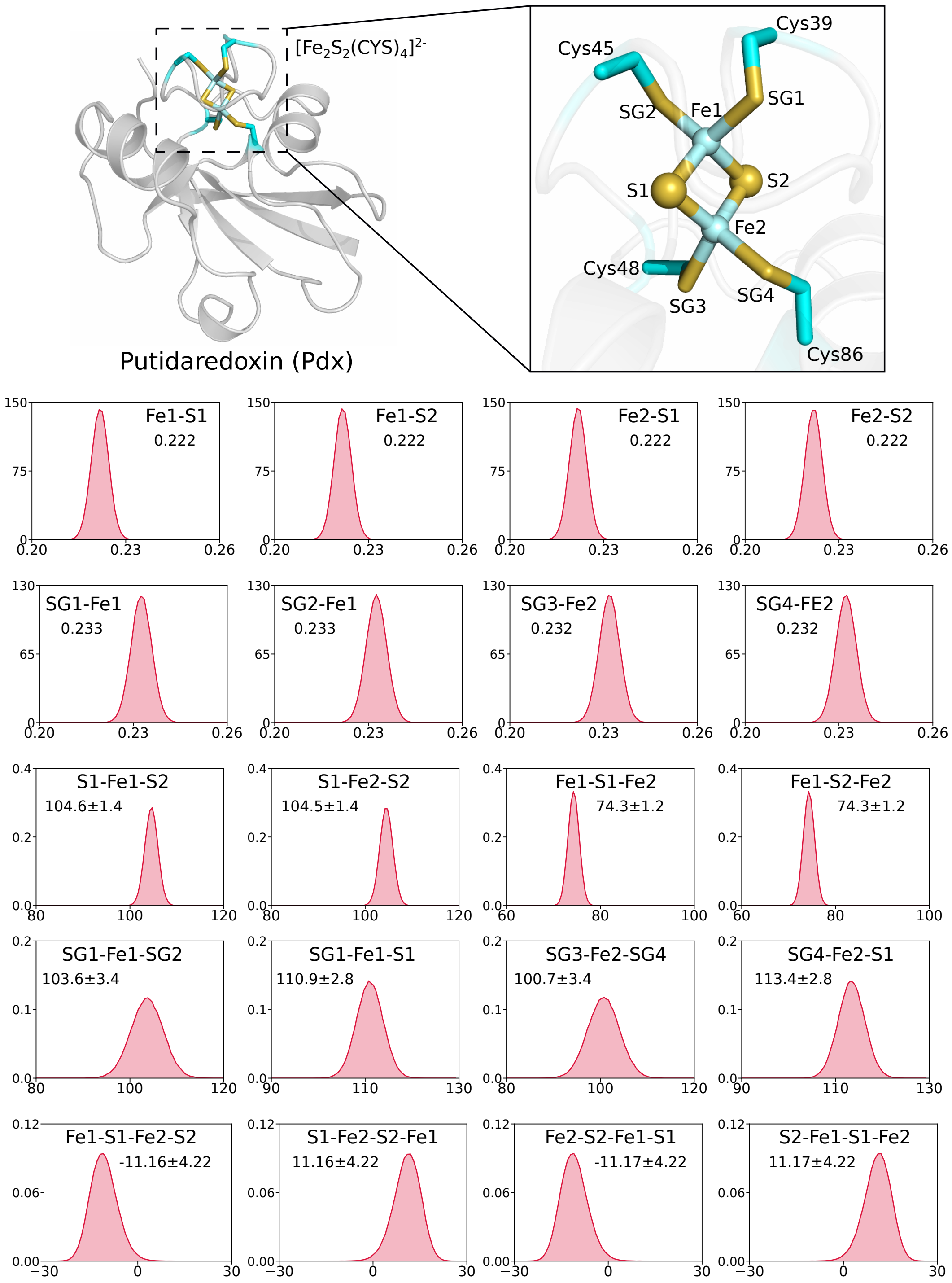


Figure S14: (Top) Crystal structure of Pdx along with constituent $Fe_2S_2$ moeity coordinated to 4 cysteine residues. Highlighted structure in inset was optimized in QM simulations. (Bottom) Bonded parameters of $[Fe_2S_2(CYS)_4]^{2-}$ as measured in large pdx simulation ensemble, depicting the stable configuration during the unbiased simulations. Each proability distribution plot is titled by atoms involved and fitted mean and standard deviation values. Plots titled by 2, 3 and 4 atoms represented bond length, angle and dihedral respectively. Bond lengths were measured in nm, angle and dihedrals were in degrees. All bond lengths had standard deviation of 0.003 nm.

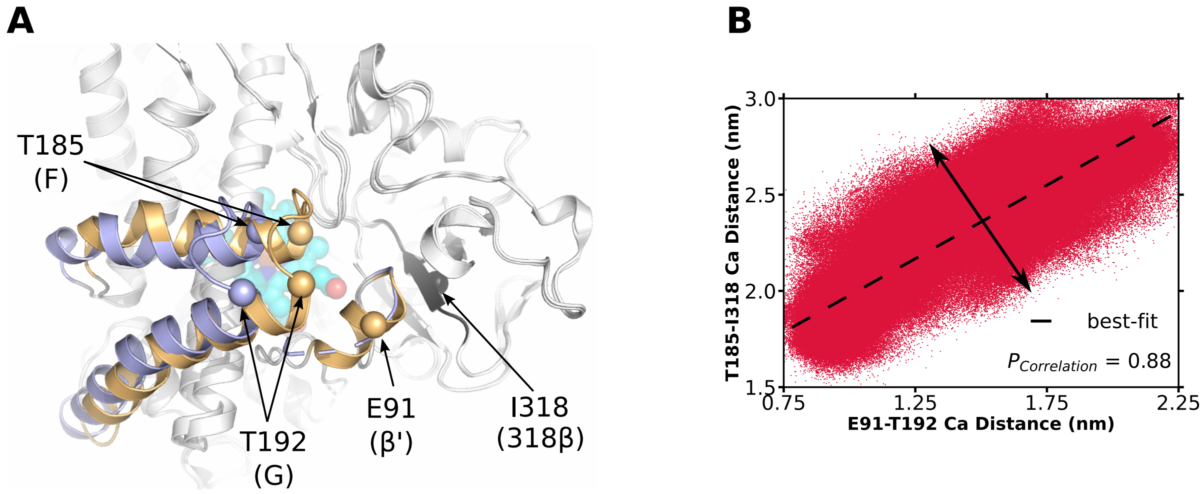


Figure S15: (A) Open (lightblue) and closed (lightorange) configurations of P450cam corresponding to PDB IDs 3L62 and 3L63 respectively. Important residues used to measure channel-1 opening were shown as spheres. (B) Correlated yet independent motions of F and G helices as observed in simulations and measured by here devised 2 distance based CVs. Although, the two CVs are linearly correlated yet to certain extent are also independent from each other. Best fit shows the correlation and spread of scatter (double headed arrow) shows the independence.

### Supplementary Results

#### SR-1: Mechanism of allosteric linkage between three binding modes of 3site state

Although the three binding modes were found to be reliably stable and reproducible (main-text), the 3site state can only be meaningful if the active and allosteric sites are allosterically linked. To understand whether the three binding modes of 3site state are functionally independent of each other or allosterically linked together, we probed the mechanism of allosteric linkage between active and allosteric sites by protein ensemble analysis approach. The systematic comparison was conducted over structural ensembles of different states of P450cam to figure out the specific conformational changes induced by the three binding modes. By comparing the different ensembles, it was evident that allosteric and active site can potentially influence each other through the correlated motions of E, F, G, H and I helices which together lies between and joined the two sites (Figure S16A). Specifically, three helical motions were found crucial in allosteric linkage between the two site; namely I helix, H helix and F/G helices.

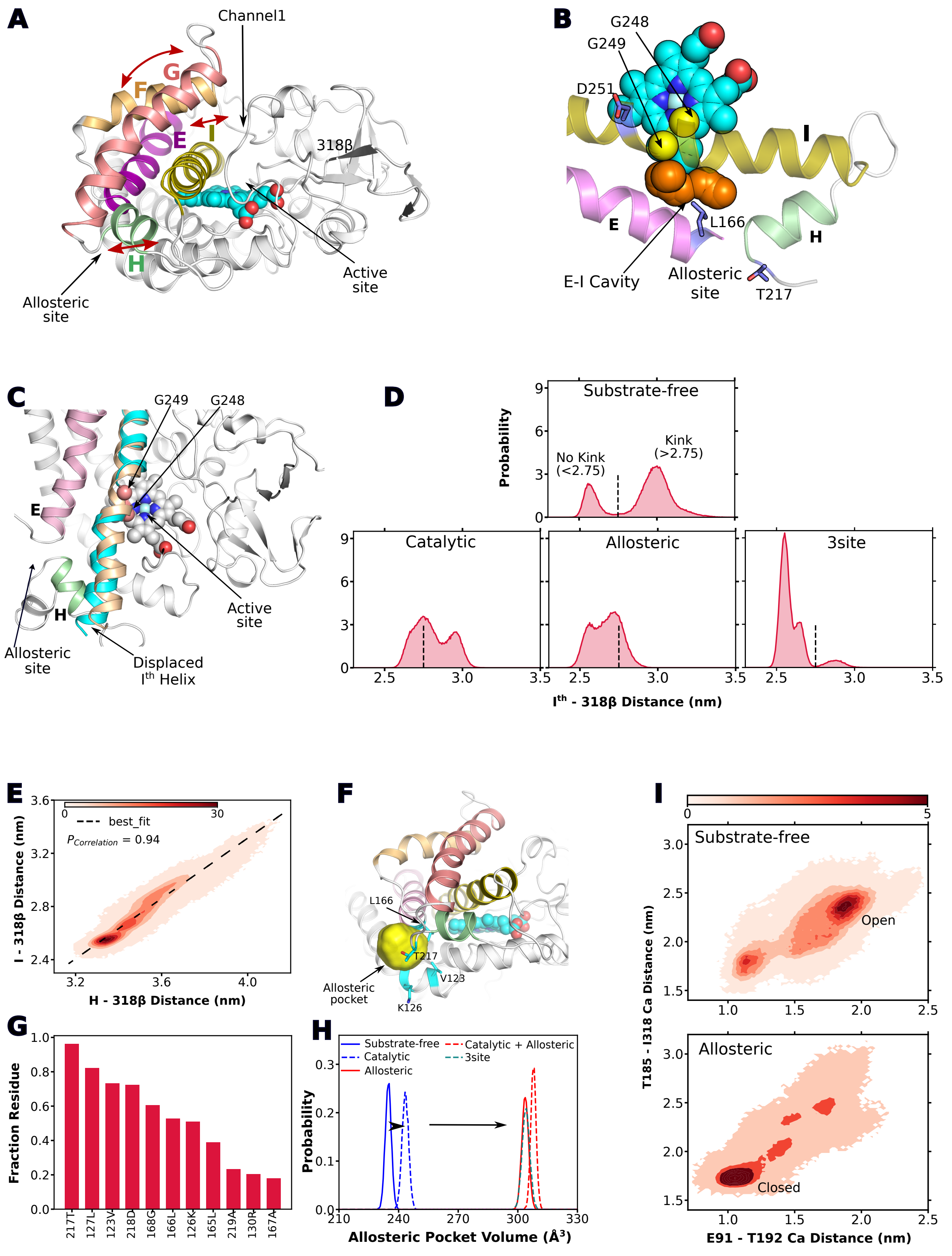


Figure S16: **Allostery in 3site state**: (A) Important secondary structures joining active and allosteric sites i.e., E, F, G, H and I helices. Arrows indicate direction of motions of the

***continued caption...*** *secondary structures, for simplicity. (B) Kink generating glycine residues in I helix and E-I cavity (orange spheres). (C) 'Kink' (cyan) and 'no-kink' (wheat) conformations of I helix. (D) State of I helix kink in different states of P450cam as measured by $I^{th}$-318β distance. Dashed line represent $I^{th}$-318β distance of 2.75 nm, cutoff used for defining 'kink' and 'no-kink' conformations. (E) Correlation between motions of I and H helices. Red surface represent the probability distribution. (F) Allosteric pocket of P450cam. Yellow surface represents the surface architecture of allosteric site. (G) Constituent residues of allosteric site. 'Fraction residue' indicate weighted portion of residue constituting the allosteric site. (H) Allosteric pocket volume in different states of P450cam. (I) Channel-1 conformation in substrate-free and allosteric simulation ensembles.*

Firstly, the most significant motions were observed for first half of I helix (residues 234-250), which possesed two significant characteristics i.e., first, the I helix was constituted by two low helical propensity and kink generating glycine residues (G248, G249); and secondly, an empty space is present nearby I helix (E-I cavity) (Figure S16B). Together, the glycine residues and E-I cavity allowed development of kink within I helix and shifting of its first half towards E and H helices or away from 318$\beta$ (Figure R1C). The flexible I helix sampled large motions that were measured with respect to stable 318$\beta$ (stable reference $\beta$-strand containing residue 318). I Helix was observed to exist in two conformations in substrate-free state i.e., *kink* where first half of I helix displaced away from 318$\beta$ and towards E and H helices or allosteric site; and *'No kink'* where I helix did not show any kink (Figure S16C,D). Upon binding in active site, the catalytic mode interacts with I helix residues and slightly reduced the kink i.e., 'no-kink' conformation (Figure S16D). Whereas allosteric mode upon binding in the allosteric site significantly pushed the I helix towards "No kink" conformation (Figure S16D).

Secondly, the H helix, constituent of allosteric site, was found to exhibit correlated motion with the I helix such that conformation of I helix can control the allosteric pocket (Figure

S16E). Shifting of I helix towards H helix ('kink' conformation) simultaneously shifted the H helix outward thereby reducing the allosteric pocket volume. Allosteric site represents a cryptic pocket formed on the back side of F and G helices and near junction of D, E and H helices consituted by residues K126, L166, T217 etc with a mean pocket volume of 234.8 $A^3$ in substrate-free state (Figure S16F-H). Binding in active site which reduced the kink in I helix increased the allosteric pocket volume to 243.4 $A^3$ whereas binding in allosteric site which significantly reduced the kink in I helix also significantly increased the allosteric pocket volume to 303.6 $A^3$ (Figure S16H). Overall, binding of allosteric mode requires enlargement of allosteric site which was found to be favoured by binding in active site.

Thirdly, significant motions were observed for F and G helices which can traverse in open and closed conformations of channel-1. As measured by T185-I318 and E92-T192 C$\alpha$ distances (methods), binding of allosteric mode near back side of F and G helices i.e., allosteric site, pulls F/G helices towards the closed conformation (not completely closed) which otherwise favoured the open conformation in substrate-free state (Figure S16I). The shifting of F and G helices towards closed conformation of channel-1 shall stabilize the active site binding modes i.e., catalytic and waiting.

Overall, binding in active site can potentially facilitate in stabilizing the allosteric mode by increasing the allosteric pocket volume via I and H helices movement, whereas binding in allosteric site can potentially stabilize active site binding modes (catalytic and waiting) by pushing on F and G helices towards closed state. Through the correlated motions of F, G, H and I helices the binding modes within active and allosteric sites can stabilize each other and hence followed the principle of allosteric reciprocity. The explicit stability calculations were provided in maintext.

#### SR-2: Allosteric mode closed channel-2

Additionally, the three binding modes also allosterically controlled the channel-2 which is the product egress route for catalytic mode (maintext). Channel-2 represented a roughly

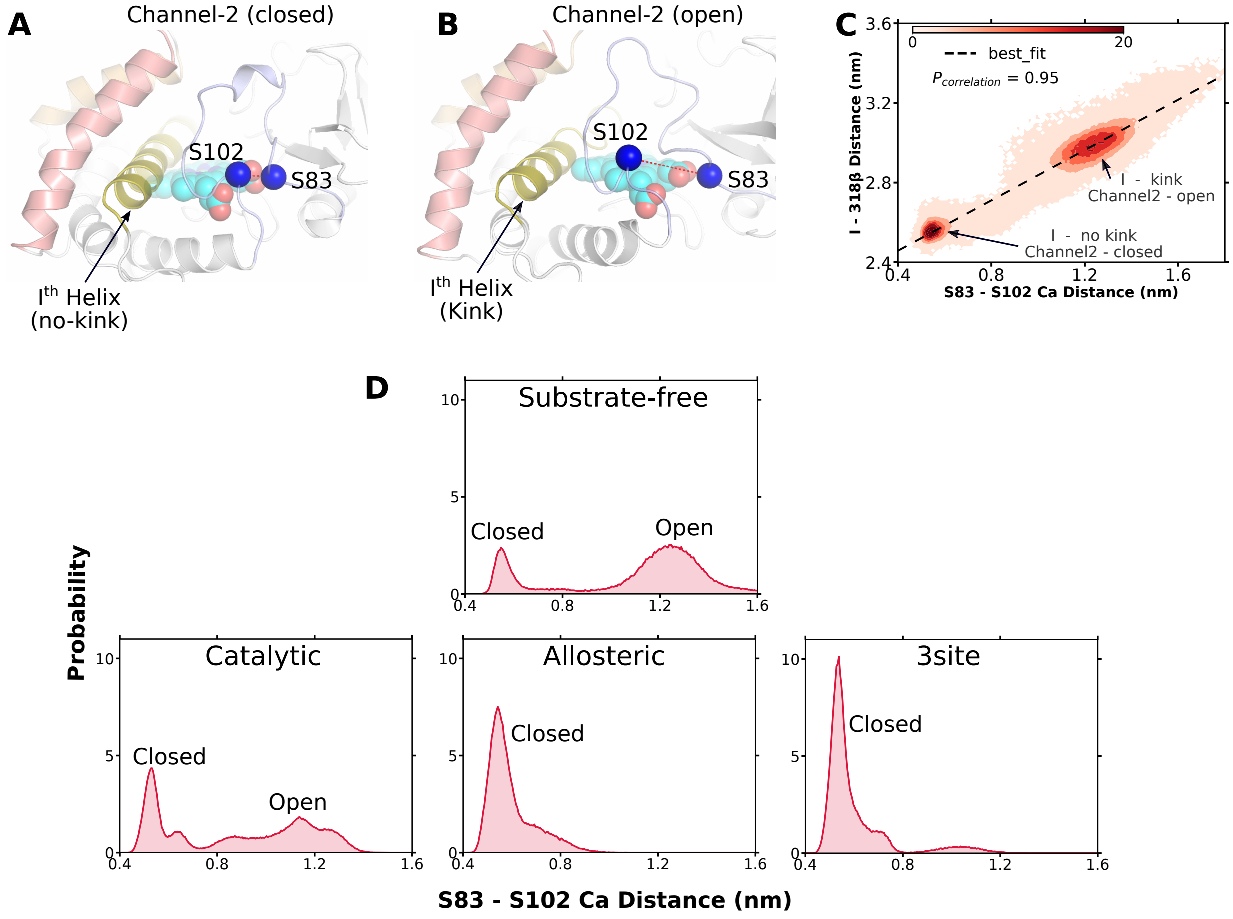


Figure S17: **Allosteric control of channel-2**: (A,B) Closed and open conformations of channel-2 in P450cam as measured by segregation of S83 and S102 C$\alpha$ atoms. State of I helix kink is also shown. (C) Correlation between channel-2 opening and kink generation in I helix. (D) Channel-2 conformation in different state of P450cam. S83-S102 C$\alpha$ distance of 0.8 nm or lower represents the closed conformation.

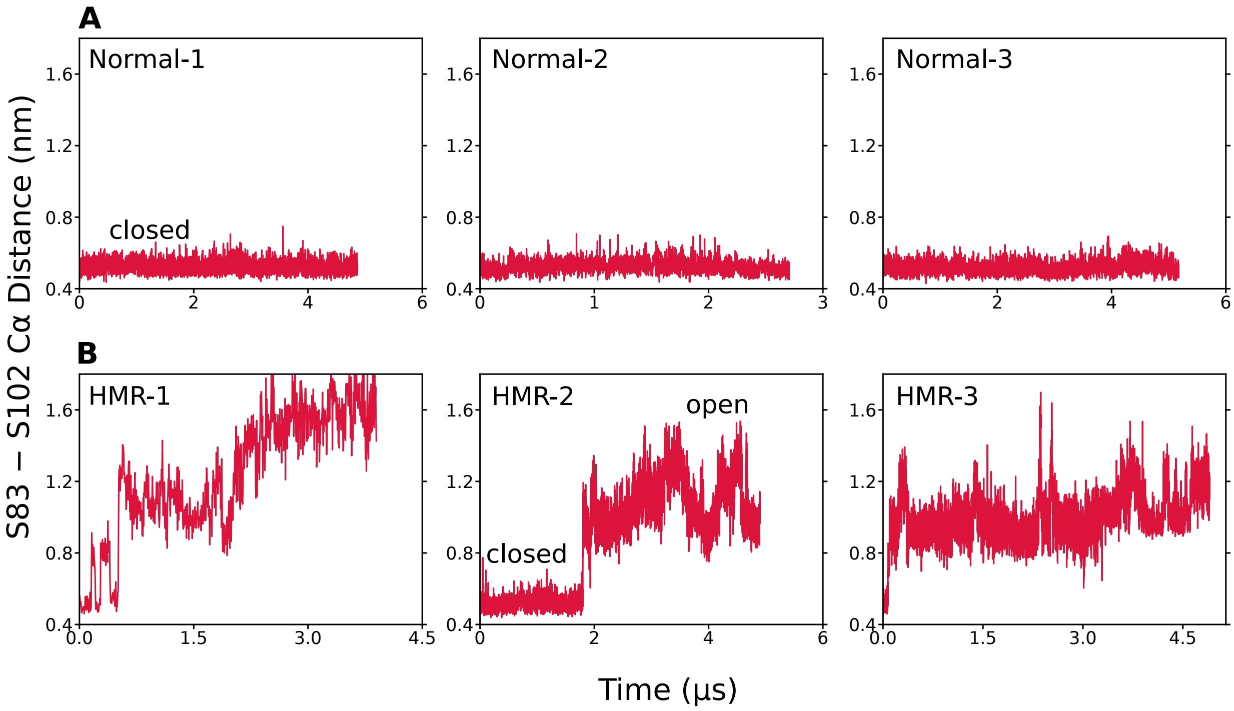


Figure S18: **Channel-2 opening in HMR simulations**: (A,B) Channel-2 conformation in normal (A) and HMR (B) binding simulations. Channel-2 spontaneously opens in HMR simulations.

orthogonal route with respect to channel-1 and lay out a direct egress path from active site to bulk solvent. Channel-2 can exist in open and closed conformation marked by distance between S83 and S102 residues such that separation of loops constituting S83 and S102 yields an opening i.e., channel-2 (Figure S17A,B). The loop constituting the S102 (i.e., 100-105 residues) govern the opening of channel-2 which in turn governed by I helix. The displacement of first half of I helix (i.e., kink conformation) allows S102 to separate from S83, hence opening the channel-2. Overall, the kink in I helix and channel-2 opening were linearly correlated (Figure S17C) such that binding modes control the I helix and inturn controlled the channel-2. Corresponding to I helix conformations, the channel-2 exists in both open and closed conformations in substrate-free state (Figure S17D). The binding in active site (catalytic) slightly shifts the equilibrium towards closed conformation while binding in allosteric site significantly induced the closing of channel-2. In 3site state, channel-2 remain predominantly in closed conformation, hence unbinding through channel-2 shall require its simultaneous opening (maintext).

Recently, allosteric site has been characterized as an additional binding site in P450cam,[27] as observed in this work also. But at odds with 3site model, allosteric mode was claimed to open up the channel-2. Although no allosteric mechanism has been elucidated in the said work,[27] we believe that the observation might be due to increased protein flexibility observed in Hydrogen Mass Repartitioned (HMR) simulations and are not the preferred choice for binding simulations.[28] Nevertheless, to explicitly confirm that channel-2 can spontaneously open up in HMR simulations, 3 binding simulations with HMR topology were performed. The system was built exactly as described for regular simulations using closed P450cam i.e., 2CPP (methods), but the topologies of protein and camphor were repartitioned according to protocol described by Hopkins et al,[29] by using a scaling factor of 3. If the normal (without HMR) simulations were performed with closed conformation of P450cam, the channel-2 was not able to open even though substrates bound to allosteric site (Figure S18A). In HMR simulations on the hand, channel-2 opened up in all the simulation trajectories (Figure S18B), clearly indicating that HMR induced increase in protein flexibility can open up channel-2. Free energy calculations using umbrella sampling approach is known to give accurate results both in normal and HMR simulations,[29] which clearly exhibit that allosteric site induced closing of channel-2 (maintext). This work is in agreement with Follmer et al,[27] where channel-2 was first claimed as egress route but opposed to the speculated dynamics of channel-2 in the said work.

#### SR-3: Decoupling the active and allosteric sites by L166A mutation

In the allosteric mechanism of P450cam, the I helix emerge out to be the most important secondary structure which governs the allosteric linkage between active and allosteric sites and also control channel-2 opening/closing transitions. Interestingly, here assigned 'kink' and 'no-kink' conformations of I helix has been observed in crystal structures (for example PDB ids - 6NBL[30] ('kink'), 3L63[26] ('no-kink')) and I helix was also known to be among the most perturbed regions of P450cam in NMR experiments. Nevertheless, the functional implications to 'kink' and 'no-kink' conformations of I helix were assigned in this work for the first time.

To further confirm the role of I helix and evaluate the possible modulations of above described allosteric model, site directed mutants were studied. As described above, the kink in the I-helix developed due to G248 and G249 and space available due to E-I cavity. Residue L166 is a major constituent of allosteric site and also lined the E-I cavity (Figure S16B, G). Residue L166, if mutated to smaller alanine shall increased the E-I cavity and allow easy kink generation in I helix. Hence, simulation ensembles of P450cam with L166A mutation were generated in substrate-free and catalytic state. As anticipated, the I helix was found to predominantly exist only in 'kink' conformation and the probability of observing 'no-kink' conformation was zero (Figure S19A). Since the kink in I helix simultaneously shifted the H helix and reduced the allosteric pocket volume, the predominant kink conformation in L166A mutant significantly reduced the allosteric pocket volume to 82.3 $\mathring{A}^3$ (as compared to

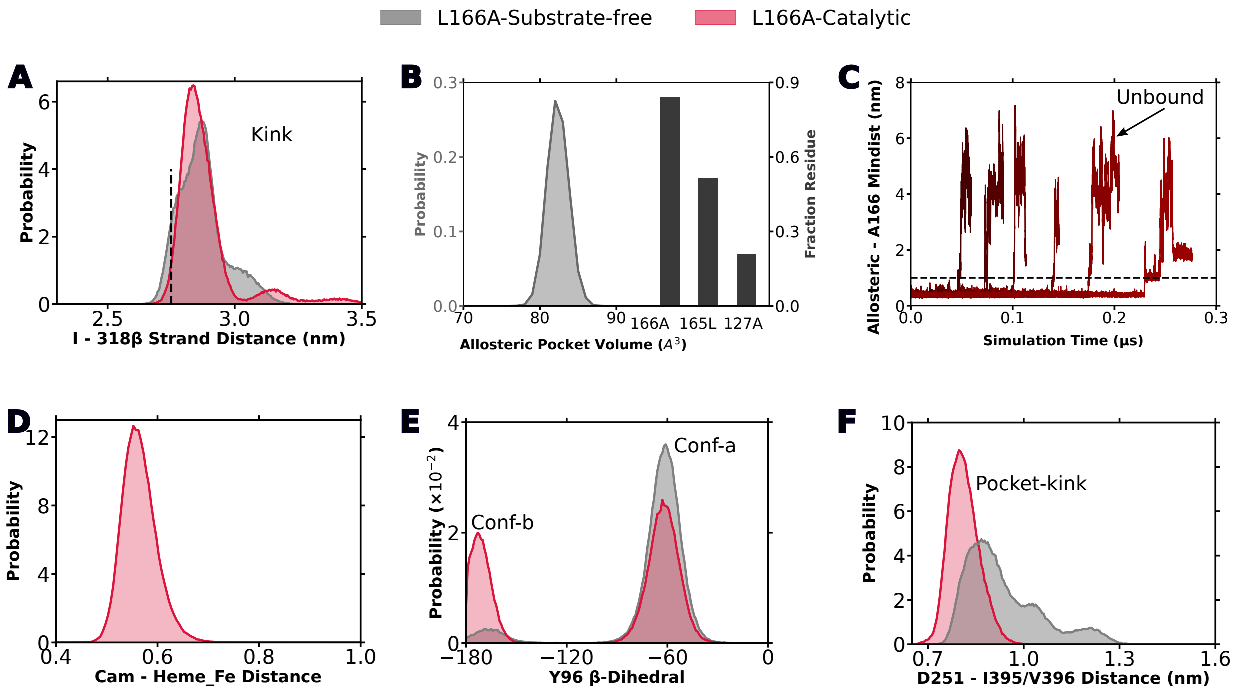


Figure S19: **Impaired allosteric site in L166A mutant**: Predominant 'kink conformation' of I helix in different states of L166A mutant P450cam. (B) Reduced allosteric pocket volume and constituent residues of allosteric site in L166A mutant. (C) Spontaneous unbinding of allosteric mode in L166A mutant within short simulation time. **Unimpaired function in active site of L166A mutant**: (D) Stable catalytic mode at cam-heme-Fe distance of 0.57 nm, in active site of L166A mutant P450cam. (E) Conformational transition of Y96 sidechain from *conf-a* to *conf-b* in active site of L166A mutant upon binding of catalytic mode, similar to wildtype (maintext). (F) pocket kink generation in active site upon binding of catalytic mode in L166A mutant, similar to wildtype.

240-300 $\AA^3$ in wildtype) constituted only by 3 residues (Figure S19B). The minute allosteric pocket of L166A was rendered incompetent to substrate binding. Simulations of L166A with camphor in allosteric site were started, it was found that substrate camphor was unable to reside in such smaller volume allosteric pocket and spontaneously unbinds within 100s of nanoseconds (Figure S19C). On the other hand, L166A did not alter the active site where catalytic mode stably reside in the active site and induced the conformational changes within the active site namely, Y96-conformation and pocket-kink (Figure S19D-F) as observed in wildtype (maintext). Overall, L166A mutation did alter the active site, instead rendered the allosteric site incompatible to binding and decoupled the active-allosteric site relationship by enhancing the kink generation in I helix.

#### SR-4: Catalytically important D251 is free of salt bridges in 3site state

D251, of I-helix and constituent of active site, is catalytically important residue which partake in reaction potentially through electron transfer such that D251 mutants are functionally inactive.[31] D251 also formed salt bridge with two residues i.e., K178 and/or R186 of F-helix.[32] F helix, along with G helix, traverse in open and closed conformations of channel-1,[26] such that in open conformation D251 is free of salt bridges; while in closed conformation, with F helix reaching nearby to I helix, D251 forms salt bridges with F helix residues.[32] Therefore based on observations till now, salt bridge status of D251 is linked with conformation of channel-1. This raised crucial question that whether the final reaction step in P450cam occurs in closed or open conformation of channel-1. NMR based observations clearly indicate that final reaction step occurs in closed conformation and in agreement with remarkably high regioselective nature of P450cam,[1] but crystallography based observations indicate that catalytically important D251 shall not partake in reaction in closed conformation of channel-1.[2]

Hereby described 3site state can explain both the substrate induced closing to channel-

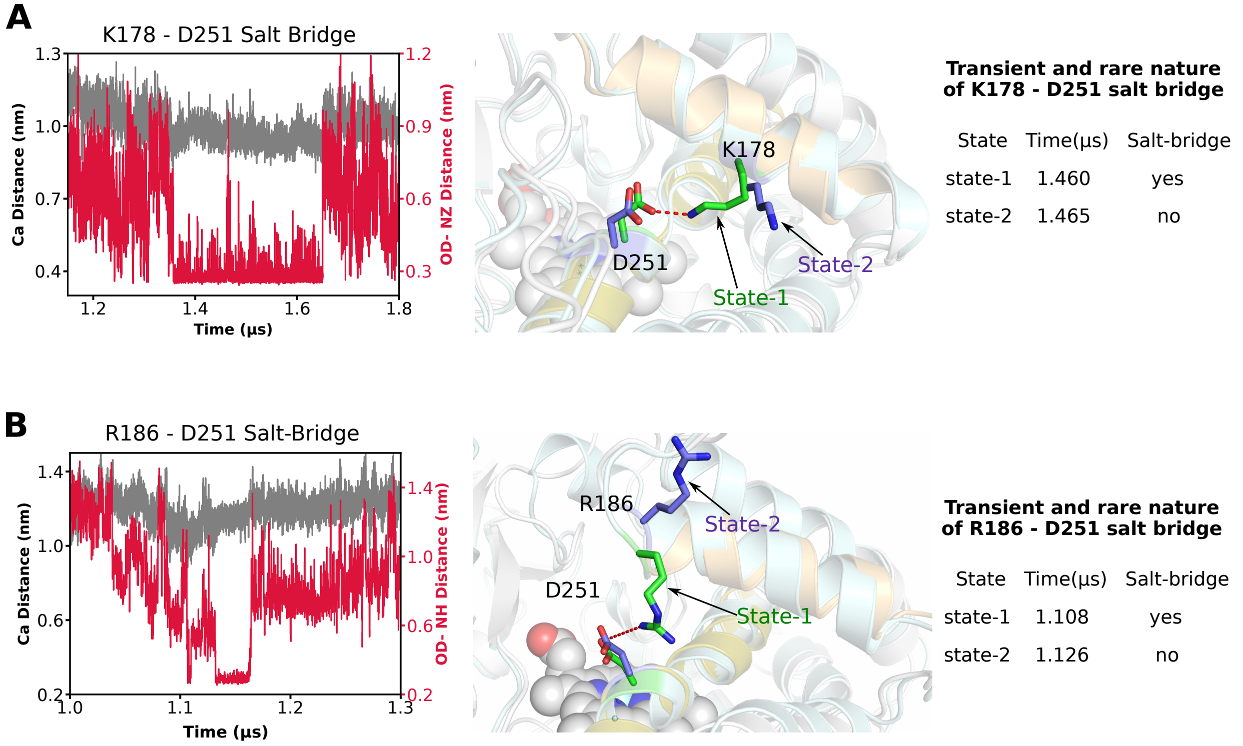


Figure S20: **Free D251 in 3site state**: (A,B) Salt bridges of D251 with K178 (A) and R186 (B) sidechains as measured by distance between terminal oxygen and nitrogen atoms of residue pairs (crimson). Values less than 4.5 Årepresents salt bridge. Gray curve represent the distance between C$\alpha$ atoms of residue pair, indicative of helical motion. Structures represent the state-1 (salt bridge) and state-2 (no salt bridge) and differed only in sidechain motions of K178 and R186. State-1 (salt bridge) attained only transiently in 3site state as shown by time analysis.

1 and conformational preference of substrate i.e., regioselectivity (maintext). Hence, 3site state, being the claimed eventual bound state of P450cam, was probed for salt bridge status of D251 with K178 and R186. A closer look at 3site state revealed that although channel-1 is in closed conformation, but D251 is largely free of its salt bridges i.e., in 3site simulation ensemble, D251 formed salt bridge with K178 and R186 only for 22% and 3% of simulation time. The transient and weak salt bridge of D251 were due to side chain motions of K178 and R186 and not due to the motions of F helix (Figure S20). Hence in 3site state, channel-1 (or F helix) sustain closed conformation while D251 remain free of salt bridges and can participate in function thereof. This indicate that the 3site state is functional. Alternatively, given that F helix is in closed conformation but D251 is free of salt bridges, the 3site state can technically be classified as 'intermediate' state also, as per recent definition of Polous et al.[33]
